# Extracellular endosulfatase Sulf-2 harbours a chondroitin/dermatan sulfate chain that modulates its enzyme activity

**DOI:** 10.1101/2021.01.04.425218

**Authors:** Rana El Masri, Amal Seffouh, Caroline Roelants, Ilham Seffouh, Evelyne Gout, Julien Pérard, Fabien Dalonneau, Kazuchika Nishitsuji, Fredrik Noborn, Mahnaz Nikpour, Göran Larson, Yoann Crétinon, Kenji Uchimura, Régis Daniel, Hugues Lortat-Jacob, Odile Filhol, Romain R. Vivès

**Author notes:** The authors contributed equally to this work (Rana El Masri, Amal Seffouh). Correspondence should be addressed to: Romain R. Vivès, IBS, 71 Avenue des Martyrs CS 10090, 38044 GRENOBLE CEDEX 9, France., and for *in vivo* studies to Odile Filhol, CEA-Grenoble, 17 Avenue des Martyrs, 38054 GRENOBLE CEDEX 9, France. **Author contributions** RElM and AS performed most of the biochemical experiments, with additional contributions from EG, FD and YC. CR and RElM performed *in vivo* experiments and data processing under the supervision of OF. JP performed SAXS analysis and modelling. KN and KU performed biochemical analysis of HSulf-2 endogenous expression. All the glycoproteomics LC-MS/MS analyses were prepared, performed and interpreted by FN, MN and GL in collaboration with the BIOMS proteomics core facility at the University of Gothenburg. RD and IS performed MS analysis. RV, OF and HLJ interpreted the data and supervised experimental work. RV, RElM, KU and OF wrote the manuscript with the help of all co-authors.

## Abstract

Sulfs represent a class of unconventional sulfatases, which differ from all other members of the sulfatase family by their structures, catalytic features and biological functions. Through their specific endosulfatase activity in extracellular milieu, Sulfs provide an original post-synthetic regulatory mechanism for heparan sulfate complex polysaccharides and have been involved in multiple physiopathological processes, including cancer. However, Sulfs remain poorly characterized enzymes, with major discrepancies regarding their *in vivo* functions. Here we show that human Sulf-2 (HSulf-2) features a unique polysaccharide post-translational modification. We identified a chondroitin/dermatan sulfate glycosaminoglycan (GAG) chain, attached to the enzyme substrate binding domain. We found that this GAG chain affects enzyme/substrate recognition and tunes HSulf-2 activity *in vitro* and *in vivo* using a mouse model of tumorigenesis and metastasis. In addition, we showed that mammalian hyaluronidase acted as a promoter of HSulf-2 activity by digesting its GAG chain. In conclusion, our results highlight HSulf-2 as a unique proteoglycan enzyme and its newly-identified GAG chain as a critical non-catalytic modulator of the enzyme activity. These findings contribute in clarifying the conflicting data on the activities of the Sulfs and introduce a new paradigm into the study of these enzymes.

## Introduction

Eukaryotic sulfatases have historically been defined as intracellular exoenzymes participating in the metabolism of a large array of sulfated substrates such as steroids, glycolipids, and glycosaminoglycan (GAGs), through hydrolysis of sulfate ester bonds under acidic conditions (Hanson et al., 2004). However, the field took a dramatic turn two decades ago, with the discovery of the Sulfs (Dhoot et al., 2001; Morimoto-Tomita et al., 2002). Unlike all other sulfatases, Sulfs were shown to be extracellular endosulfatases that catalyzed the specific 6-*O*-desulfation of cell-surface and extracellular matrix heparan sulfate (HS), a polysaccharide with vast protein binding properties and biological functions (El Masri et al., 2017; Li and Kusche-Gullberg, 2016; Sarrazin et al., 2011). And unlike all other sulfatases, Sulfs could not be related to a straightforward metabolic function, but rapidly emerged as a novel major regulatory mechanism of HS biological activities, with roles in many physiopathological processes, including embryonic development, tissue homeostasis and cancer (Bret et al., 2011; Rosen and Lemjabbar-Alaoui, 2010; Vives et al., 2014).

Sulfs share a common molecular organization (Figure S1). The furin-processed mature form features a general sulfatase-conserved N-terminal catalytic domain (CAT) including the enzyme active site (and notably, the catalytic *N*-formylglycine (FGly) converted cysteine residue), and a unique highly basic hydrophilic domain (HD), which shares no homology with any other known protein and is responsible for high affinity binding to HS substrates (Ai et al., 2006; Frese et al., 2009; Seffouh et al., 2013, 2019a; Tang and Rosen, 2009). Sulfs display a number of post-translational modifications (PTM)(Morimoto-Tomita et al., 2002). Furin cleavage (Tang and Rosen, 2009) and *N*-glycosylations (Ambasta et al., 2007; Seffouh et al., 2019b) may be dispensable for the enzyme activity, but play a role in the enzyme attachment to the cell surface, while conversion of C_88_ into a FGly residue is a hallmark of all sulfatases and is essential for the catalytic activity (Dierks et al., 2005). Recent studies reported that human Sulfs (HSulfs) catalyzed the 6-*O*-desulfation of HS through an original, processive and orientated mechanism (Seffouh et al., 2013), and that substrate recognition by the enzyme HD domain involved multiple, highly dynamic, non-conventional interactions (Harder et al., 2015; Walhorn et al., 2018).

However, despite increasing interest, Sulfs remain to be highly elusive enzymes. Little is known about their molecular structures, catalytic mechanisms and substrate specificities. Our limited understanding of these enzymes is well illustrated by the wealth of conflicting data in the literature, reporting major discrepancies between *in vitro* and *in vivo* data, according to the biological system or the enzyme isoforms considered. This is particularly clear in cancer, where both anti-oncogenic and pro-oncogenic effects of the Sulfs have been reported (Rosen and Lemjabbar-Alaoui, 2010; Vives et al., 2014; Yang et al., 2011).

## Results

### HSulf-2 is an enzyme modified with a CS/DS chain

Recently, we achieved for the first time high yield expression and purification of HSulf-2, which paved the way to progress in the biochemical characterization of this enzyme (Seffouh et al., 2019a). Surprisingly, the purification step of size-exclusion chromatography highlighted an unexpectedly high apparent molecular weight (*aMW*) for the enzyme (> ~1000 kDa, for a theoretical molecular weight of 98170 Da, Figure 1A), although possible protein aggregation or high order oligomerization were ruled out by quality control negative staining electron microscopy (Seffouh et al., 2019a). Noteworthy, we also failed to detect the C-terminal chain containing the enzyme HD domain using PAGE analysis (Figure 1D, lane 1), even if the presence of both chains was ascertained by Edman N-terminal sequencing (Seffouh et al., 2019a). In line with this, we previously reported unusually weak mass spectrometry ionization efficiency of the HSulf-2 C-terminal chain (Seffouh et al., 2019b). Small angle X-ray scattering (SAXS) analysis of the protein yielded Guinier plots and pair-distribution function in accordance with a Dmax of 40+/-3 nm, suggesting an elongated molecular shape with an *aMW* of ~700 kDa, which supported our size-exclusion chromatography data (Figure S2A-E). Furthermore, results suggested the presence of two distinct domains within HSulf-2: a globular domain and an extended, flexible, probably partially unfolded region. Interestingly, similar analysis performed on a HD-devoid HSulf-2 variant (HSulf-2 ΔHD) showed only the globular domain (Figure S2F-K), which 11 nm size fitted that of a modelled structure of the CAT domain (Figure S2K). However, it seemed unlikely that the HD domain on its own could account for the second, large flexible region. We thus initially speculated that the enzyme could have been purified in complex with HS substrate polysaccharide chains. To test this, HSulf-2 was treated with heparinases (to digest potentially bound HS substrate) or with chondroitinase ABC (to digest non-substrate GAGs of CS/DS types) prior to size-exclusion chromatography. Results showed no effect of the heparinase treatment (Figure S3B), while digestion with chondroitinase ABC dramatically delayed HSulf-2 size-exclusion chromatography elution time, thus indicating the presence of CS/DS associated to the enzyme (Figure 1B). Attempts to dissociate the HSulf-2-CS/DS complex with high NaCl concentrations were unsuccessful (Figure S3C), thereby suggesting covalent linkage between the polysaccharide and the protein. In addition, chondroitinase ABC treatment allowed for detection of a broad additional band of ~50 kDa apparent molecular weight (Figure 1D). This band was assigned to the enzyme C-terminal subunit, as confirmed by Western blot (WB) analysis (Figure 1D). Of note, the HSulf-2 ΔHD variant (lacking the HD but not the enzyme C-terminal region) did not exhibit such band on PAGE (Seffouh et al., 2019a). We therefore concluded from these results that a CS/DS chain was covalently attached to the HSulf-2 HD domain. The presence of such a chain could account for the high *aMW* determined by SAXS and size-exclusion chromatography, and could also hinder migration/detection of the C-terminal subunit in PAGE/WB.

**Fig 1:**
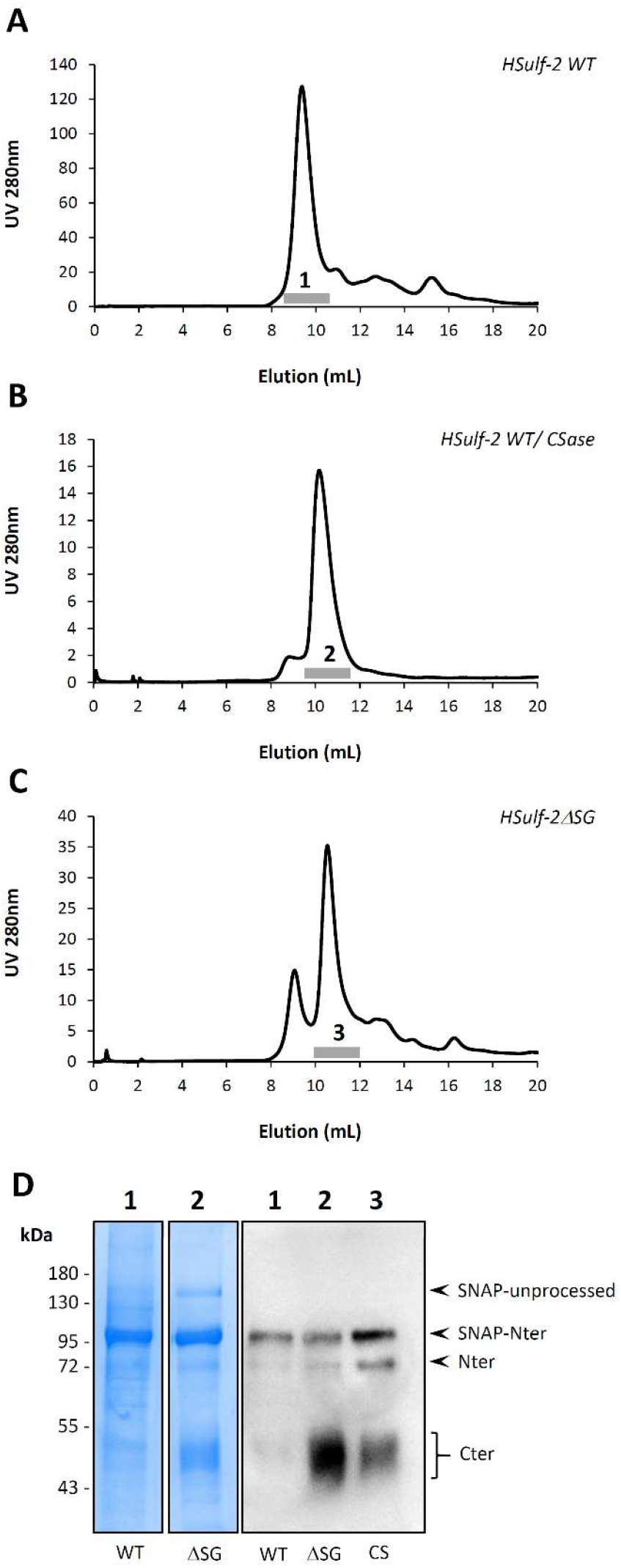
Purification and characterization of HSulf-2 and HSulf-2ΔSG. Size exclusion chromatography profile of HSulf-2 WT (**A**), chondroitinase ABC pre-treated HSulf-2 (**B**) and HSulf-2ΔSG (**C**); grey bars indicate Sulf-containing fractions. (**D**) PAGE/Coomassie blue staining and Western blot analysis of HSulf-2 WT (WT, lanes 1), HSulf-2ΔSG (ΔSG, lanes 2) and chondroitinase ABC pre-treated HSulf-2 (CS, lane 3), using the anti HD H19 antibody. Analysis shows a ~95 kDa band corresponding to HSulf-2 N-terminal subunit in fusion with the SNAP-tag (SNAP-Nter) and multiple/broad ~50 kDa bands corresponding to the C-terminal subunit, which includes HSulf-2 HD domain (Cter). Of note, a residual ~75 kDa band corresponding to the N-terminal subunit lacking its SNAP-tag could also be detected (Nter). In addition, Coomassie blue staining but not WB, revealed the presence of a full-length, unprocessed GAG-free HSulf-2 form (SNAP-unprocessed).

GAGs are covalently bound to specific glycoproteins termed proteoglycans (PGs), through a specific attachment site involving the serine residue of a SG dipeptide, primed by a xylose residue (Esko and Zhang, 1996). Xylosides are widely used inhibitors of GAG assembly on such motifs (Chua and Kuberan, 2017). As such, size-exclusion chromatography analysis of HSulf-2 expressed in xyloside-treated HEK 293 cells showed a dramatic reduction of the high *aMW* form and concomitant increase of a form eluting as the chondroitinase ABC-treated HSulf-2 (Figure S3D), further supporting the presence of a covalently attached GAG chain. Examination of HSulf-2 amino-acid sequence showed two SG dipeptides: S_508_G and S_583_G. We thus expressed and produced a HSulf-2 variant lacking these two motifs (HSulf-2ΔSG). The HSulf-2ΔSG variant eluted at the same time as seen in the chondroitinase ABC-treated wild type (WT) HSulf-2 (Figure 1C), with restored detection of the C-terminal chain by Coomassie blue-stained PAGE and WB analysis (Figure 1D). Both SG dipeptides are located within the enzyme HD domain, but on each side of the furin cleavage site (Figure S1). As our PAGE/WB data located the CS/DS chain on the C-terminal subunit, we thus speculated that the S_583_G motif downstream the furin cleavage sites was the actual GAG attachment site on HSulf-2. To assess this, we performed single mutations of the first and second sites. Size-exclusion analysis of the resulting variants validated the presence of a CS/DS-type GAG chain on the S_583_G, but not S_508_G, dipeptide motif (Figure S3E and S3F). Finally, we also confirmed that the presence of N-and C-terminal tags did not bias the results, as tobacco etch virus (TEV) protease digestion did not affect size-exclusion chromatography elution times of HSulf-2 WT, chondroitinase ABC-treated HSulf-2 WT or HSulf-2ΔSG (Figure S4).

To characterize the HSulf-2 GAG chain further, we analyzed both HSulf-2 WT and HSulf-2ΔSG variants by mass spectrometry. MALDI-TOF MS analysis of HSulf-2ΔSG showed a major peak at *m/z* 53,885 that we assigned to the doubly charged ion [M+2H]^2+^ of the whole HSulf-2 variant. Interestingly, corresponding mono- and triple-charged ions at *m/z* 108,250 and 37,180 were also observed. Based on this distribution of multiple charged ions, an average experimental mass value of 108,388 ± 506 g.mol^-1^ was determined for the whole HSulf-2ΔSG. HSulf-2ΔSG thus exhibited a ~25,000 Da lower average mass value compared to HSulf-2 WT average molecular weight previously determined by MALDI MS (133,115 Da) (Seffouh et al., 2019b). Such mass difference likely corresponds to mass contribution of the GAG chain. (Figure S5).

Covalent linkage of a ~25 kDa sulfated GAG polysaccharide to HSulf-2 would result in a huge increase of the hydrodynamic volume, which is consistent with the high *aMW* observed in size-exclusion chromatography (Figure 1A) and in SAXS analysis (Figure S2). Altogether, these data provide converging evidence that HSulf-2 features a unique PTM at the level of its HD domain, corresponding to a covalently-linked CS/DS polysaccharide chain. This result thus highlights HSulf-2 as a new member of the large PG family.

### Endogenous expression of GAG-modified HSulf-2

We Identified a GAG chain on HSulf-2 when overexpressed in HEK transfected cells. To confirm the physiological relevance of these findings, we sought to demonstrate the presence of this GAG chain on the naturally occurring enzyme. In that attempt, we first used a strategy originally designed to identify new proteoglycans (Noborn et al., 2015). Nano-scale liquid chromatography MS/MS analysis of trypsin- and chondroitinase ABC-digested PGs isolated from the culture medium of human neuroblastoma SH-SY5Y cells led to the identification of a HSulf-2 specific, 21 amino acid long glycopeptide highlighting a CS/DS attachment site on the S_583_ residue of HSulf-2 (Figure S6).

To get further insights into GAG modification of endogenously expressed HSulf-2, we analyzed HSulf-2 expressed by two additional cell types: MCF7 human breast cancer cells and human umbilical vein endothelial cells (HUVECs). Detection and characterization of endogenous Sulfs were challenging. Expression yields are usually low, and WB immunodetection yields different band patterns, depending on cells, PTMs and furin cleavages (see Figure S1). To address these issues, we made use of Sulf high *N*-glycosylation content and used a protocol of culture medium enrichment based on a lectin affinity chromatography. We analyzed enriched conditioned medium by WB, using antibodies raised against either HSulf-2 N-terminal (H2.3) (Uchimura et al., 2006) or C-terminal (2B4) (Lemjabbar-Alaoui et al., 2010) subunits (Figure S1). WB analysis of HSulf-2 secreted in the conditioned medium of MCF7 using 2B4 yielded broad diffuse bands, respectively in the ~66-130 and ~185-270 kDa size range (Figure 2A, lane 1). We attributed these bands to CS/DS-conjugated C-terminal subunit fragments and to a CS/DS-conjugated full-length, furin-uncleaved HSulf-2 form, respectively. The presence of the CS/DS chain was confirmed by chondroitinase ABC treatment, which converted the broad bands into two sets of well-defined bands, corresponding to GAG-depolymerized C-terminal fragments and full-length unprocessed forms (Figure 2A, lane 2). These changes could definitely be attributed to the cleavage of the GAG chains, as the treatment with heat inactivated chondroitinase ABC showed the same band pattern as the non-treated samples (Figure 2A, lane 3).

**Fig 2:**
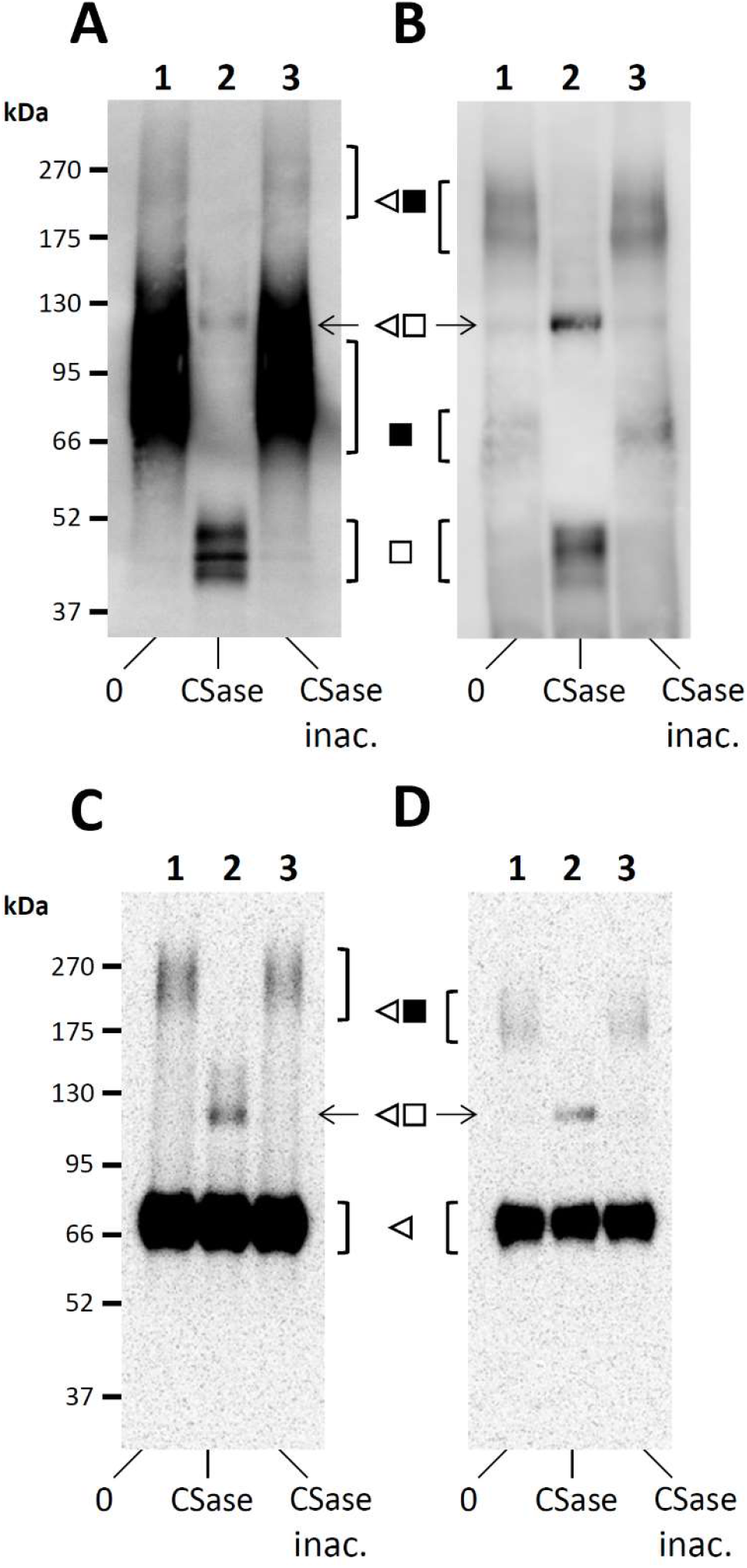
Endogenous expression of GAG bearing HSulf-2 in MCF7 and HUVEC cells. Western blot analysis of pre-purified concentrated conditioned medium from MCF7 (**A, C**) and HUVEC (**B, D**) using anti C-ter 2B4 (**A, B**) and anti N-ter H2.3 antibodies (**C, D**), prior to (0, lanes 1) or after treatment with chondroitinase ABC (CS, lanes 2). Digestions with heat-inactivated chondroitinase ABC were used as controls (CS inac., lanes 3). The nature of detected bands is shown as follow: black symbols for CS/DS conjugated fragments; white symbols for GAG-free fragments; triangles for the N-terminal subunit, squares for the C-terminal subunit. Of note, analysis indicate the presence in both samples of unprocessed forms (triangle + square, sharp band at ~125 kDa), and at least in the HUVEC conditioned medium, the presence of GAG-free HSulf-2 forms (bands corresponding to C-ter fragments within the 40-57 kDa MW range, and an unprocessed form at 125 kDa detected in the untreated samples, gel B lanes 1 and 3).

Analysis of HSulf-2 from HUVEC pre-purified conditioned medium yielded similar results (Figure 2B), but with remarkable differences. First, detected signals were markedly less intense. Although quantification by immunodetection should be cautiously considered, this suggested that expression levels of HSulf-2 were different between MCF7 and HUVEC cells. We also noticed discrepancies regarding furin processing activity (distinct ratios between processed and unprocessed forms). WB analysis using the N-terminal reactive H2.3 antibody confirmed the presence of the N-terminal subunit being unaffected by the chondroitinase ABC treatment (~75 kDa size), and supported the identification of full-length unprocessed forms within the analyzed samples (~160-250 kDa size range, Figure 2C and 2D). Finally, although WB analysis showed similar band patterns for these two cell lines (Figure 2A and 2B), GAG-conjugated fragments from HUVEC cells migrated at slightly lower *aMW* (~60-75 and ~160-250 kDa, Figure 2B, lane 1). In addition, we also detected bands corresponding to GAG-lacking fragments, for HSulf-2 from HUVEC at least (C-ter and unprocessed enzyme, see Figure 2C and 2D).

Altogether, these results confirm that endogenously expressed HSulf-2 harbor a CS/DS chain and indicate the co-existence of GAG-modified and GAG-free forms. Furthermore, our data suggest cell-dependent specificities of HSulf-2 PTM (e.g. furin processing, and the GAG structure), which could provide additional regulation/diversity of the enzyme structural and functional features.

### HSulf-2 GAG chain modulates enzyme activity *in vitro*

The HD is a major functional domain of the Sulfs. This domain is required for the enzyme high affinity binding to HS substrates and for processive 6-*O*-endosulfatase activity (Frese et al., 2009; Seffouh et al., 2013; Tang and Rosen, 2009). We thus anticipated that the presence of a GAG chain on this domain would significantly affect the enzyme substrate recognition and activity.

To study this, we assessed HSulf-2 WT and HSulf-2ΔSG 6-*O*-endosulfatase activities, using heparin as a surrogate of HS. We analyzed the disaccharide composition of HSulf-2 treated heparin and measured the content of [UA(2S)-GlcNS(6S)] trisulfated disaccharide, which is the enzyme’s primary substrate (Frese et al., 2009; Pempe et al., 2012; Seffouh et al., 2013). Results showed enhanced digestion of the disaccharide substrate with HSulf-2ΔSG versus HSulf-2 WT, and a concomitant increase in the [UA(2S)-GlcNS] disaccharide product (Figure 3A). We speculated that the observed increase in endosulfatase activity could be due to an improved enzyme-substrate interaction. We thus analyzed the binding of HSulf-2 WT and HSulf-2ΔSG to surface coated biotinylated heparin. Results showed an increase, although modest in HSulf-2ΔSG binding to heparin, with calculated *K*_D_s of 10.2±0.4 nM and 7.1±0.8 nM for HSulf-2 WT and HSulf-2ΔSG, respectively (Figure 3B).

**Fig 3:**
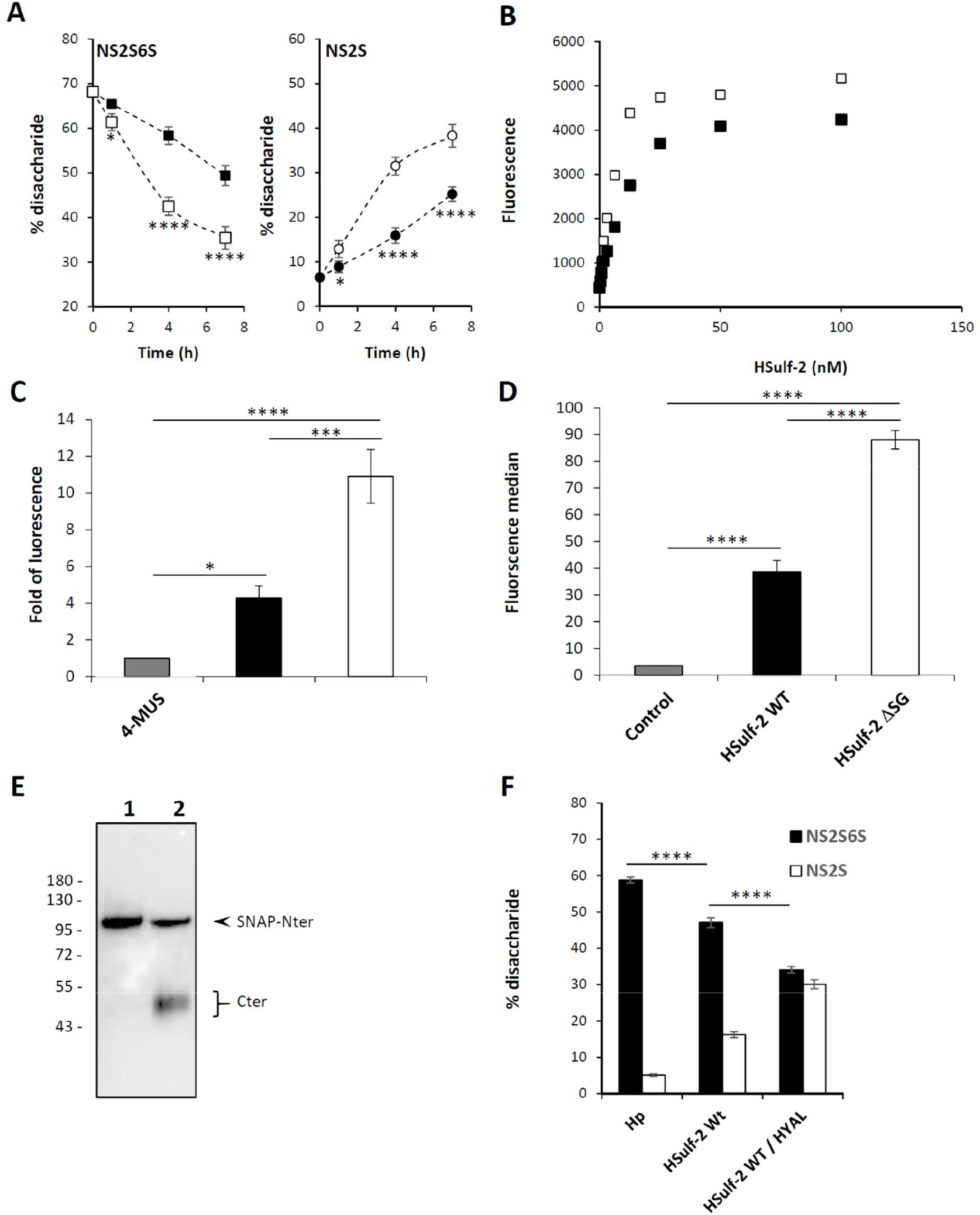
Biological activities of HSulf-2 WT and GAG-free HSulf-2. **(A)** HS 6-*O*-endosulfatase activity of HSulf-2 WT (black symbols) and HSulf-2ΔSG (white symbols) was assessed by monitoring the time course digestion of [UA(2S)-GlcNS(6S)] trisulfated disaccharides (NS2S6S, square) into UA(2S)-GlcNS] disulfated disaccharides (NS2S, circle). Data are expressed as a percentage of total disaccharide content. Ordinary two-way ANOVA with time of incubation and type of HSulf-2 as factors revealed significant effects on 6-*O*-endosulfatase activity (time: F_3, 16_ = 272.7, *P* < 0.0001; HSulf-2 type: F_1, 16_ = 140.5, *P* < 0.0001; interaction: F_3, 16_ = 28.57, *P* < 0.0001) (left panel) and a concomitant increase in the digested product (time: F_3, 16_ = 394, *P* < 0.0001; HSulf-2 type: F_1, 16_ = 195, *P* < 0.0001; interaction: F_3, 16_ = 39.94, *P* < 0.0001) (right panel). Post-hoc Bonferroni’s test showed significant difference in the HS 6-*O*-endosulfatase activity at 1h incubation and thereafter until 7h in HSulf-2 ΔSG (n=3) compared with HSulf-2 WT (n=3). Error bars indicate SD. (**B**) Binding immunoassay of HSulf-2 WT (black) and HSulf-2ΔSG (white) to a streptavidin-immobilized heparin surface. Data are representative of three independent experiments. (**C**) The aryl-sulfatase activity of HSulf-2 WT (n=3, black) and HSulf-2ΔSG (n=3, white) was measured using 4MUS fluorogenic pseudo-substrate. Results are expressed as a fluorescence fold increase compared to negative control (4MUS alone, n=3, grey). (**D**) Binding of HSulf-2 WT (n=3, black) and HSulf-2ΔSG (n=3, white) to the surface of human amnion-derived Wish cells was monitored by FACS using the H2.3 anti-HSulf-2 antibody. Ordinary one-way ANOVA with type of HSulf-2 as a factor revealed significant effects on a sulfatase activity (F_2, 6_ = 90.18, *P* < 0.0001) (**C**), and a cell-surface binding (F_2, 6_ = 536.7, *P* < 0.0001). (**D**) Post-hoc Tukey’s range test showed significant difference in the 4MUS activity and binding to human Wish cells in HSulf-2ΔSG compared with control (n=3, grey) or HSulf-2 WT. Error bars indicate SD. (**E**) Western blot analysis of HSulf-2 WT (lane 1) and Hyaluronidase treated HSulf-2 WT (lane 2), using the anti HD H19 antibody. (**F**) [UA(2S)-GlcNS(6S)] trisulfated disaccharide (NS2S6S, black) and UA(2S)-GlcNS] disulfated disaccharid (NS2S, white) content (as in (A), expressed as a percentage of total disaccharide content, n=3) of heparin, without (Hp) or after digestion with HSulf-2 WT or Hyaluronidase (HYAL)-treated HSulf-2 WT (4 h at 37 °C). Data show significantly increased heparin 6-*O*-desulfation for HYAL-treated HSulf-2 WT. Error bars indicate SD (****P<0.0001).

The second functional domain of the Sulfs, CAT, comprises the enzyme active site. CAT alone is unable to catalyze HS 6-*O*-desulfation, but it exhibits a generic arylsulfatase activity that can be measured using the fluorogenic pseudosubstrate 4-methyl umbelliferyl sulfate (4-MUS). Surprisingly, HSulf-2ΔSG showed greater (~2.5 fold increase) arylsulfatase activity than that of HSulf-2 WT (Figure 3C). We thus concluded from these observations that newly identified CS/DS chain of HSulf-2 regulates the enzyme activity, both by modulating HD domain/substrate interaction and by hindering access to the active site. We hypothesize that these effects could be due to electrostatic hindrance preventing the interaction of the enzyme functional domains with sulfated substrates.

Aside enzyme activity, the interaction of HS with the Sulf HD domain is also involved in the retention of the enzyme at the cell surface, a mechanism that may also govern diffusion and bioavailability of the enzyme within tissues (Frese et al., 2009). To investigate this, we analyzed the interaction of HSulf-2 WT and HSulf-2ΔSG with cellular HS by FACS, using human amniotic epithelial Wish cells as a model. Again, results showed a significant increase in binding of the HSulf-2 form lacking the CS/DS chain to Wish cells (Figure 3D). These data therefore suggest that HSulf-2 GAG chain may also influence enzyme retention at the cell surface.

As GAG-lacking HSulf-2ΔSG variant exhibited enhanced HS 6-*O*-endosulfatase activity, we sought to investigate whether enzymatic removal of HSulf-2 GAG chain would lead to a similar effect. Hyaluronidases are the only mammalian enzymes to exhibit chondroitinase activity (Csoka et al., 2001; Bilong M. et al., manuscript in revision). We found that treatment of Hsulf-2 WT with hyaluronidase allowed WB detection of the ~50 kDa band corresponding to the enzyme C-terminal subunit (Figure 3E), and boosted heparin 6-*O*-desulfation (Figure 3F), with an efficiency similar to that of the HSulf-2ΔSG variant.

### HSulf-2 GAG chain modulates tumor growth and metastasis *in vivo*

We next investigated the effect of HSulf-2 GAG chain on tumor progression *in vivo*, using a mouse xenograft model of tumorigenesis and metastasis. For this, we overexpressed by lentiviral transduction either HSulf-2 WT or HSulf-2ΔSG in MDA-MB-231, a human breast cancer cell line that does not express any HSulfs endogenously (Peterson et al., 2010). After selection, stable expression of Sulfs in transduced cells was confirmed by WB. In contrast, cells transduced with an unrelated protein (DsRed) showed no endogenous expression of the Sulfs (Figure S7A).

We also validated the endosulfatase activity of HSulf-2 produced in MDA-MB-231, by treating heparin with concentrated conditioned medium from transduced cells. Results showed no activity for the medium of DsRed-transfected cells, while conditioned medium from either HSulf-2 WT or HSulf-2ΔSG transduced cells efficiently digested heparin, as shown by the increase of [UA(2S)-GlcNS] disaccharide product (Figure S7B). Again, results suggested higher endosulfatase activity for HSulf-2ΔSG transduced cells. Finally, we confirmed the presence of the CS/DS chain on MDA-MB-231 HSulf-2 WT, by treating conditioned medium with chondroitinase ABC, followed by WB analysis (Figure S7C). Of note, results also showed a significant proportion of GAG-free and full-length, unprocessed forms of HSulf-2 in the chondroitinase ABC-untreated conditioned medium (Figure S7C, lane 1).

DsRed, HSulf-2 WT or HSulf-2ΔSG transduced MDA-MB-231 cells were then xenografted into the mammary gland of mice with severe combined immunodeficiency (SCID). Tumor volumes were monitored every 2 days and xenografted SCID mice were euthanized when the first tumors reached 1 cm^3^ in size (Day 52), in accordance with the European ethical rule on animal experimentation. Primary tumors, along with lymph nodes and lungs, were collected for further analysis. Results showed little effects of HSulf-2 WT expression on the tumor size (Figure 4A). Our data are therefore in disagreement with previous work, which reported either anti-oncogenic (Peterson et al., 2010) or pro-oncogenic (Zhu et al., 2016) effects of HSulf-2 WT expression in MDA-MB-231 cells using similar *in vivo* mouse models. However, it should be noted that a major difference between these three studies is the size of xenograft tumors achieved (~0.04 cm^3^ and 3-4 cm^3^ respectively, in the studies mentioned above). Such conflicting data clearly exemplify the complexity of HSulf regulatory functions and possible bias, which could result from the experimental design. In contrast, expression of the HSulf-2ΔSG variant significantly promoted tumor growth, in comparison to both DsRed and HSulf-2 WT conditions. Noteworthy, WB analysis of tumors confirmed sustained expression of the enzyme in both HSulf-2 WT and HSulf-2ΔSG - but not DsRed-conditions (Supp. Figure S7D). Histological analysis of tumor sections using an eosin/hematoxylin staining showed greater necrotic area in control tumors than in HSulf-expressing tumors (Figure 4B). As necrosis is a hallmark of hypoxia in growing tumors that is mainly due to lack of angiogenesis, we studied tumor vascularization using α Smooth Muscle Actin (αSMA) immunostaining. Results showed no apparent changes in αSMA labelling upon HSulf-2 WT expression. However, tumor vascularization was increased in HSulf-2ΔSG tumors (Figure S8A and S8B). We next analyzed lymph nodes and lungs for secondary tumors. Lung, which is a primary target for metastasis in this tumor model, was affected in all conditions (Figure 4C). However, size of metastasis-induced secondary tumors was significantly greater in HSulf-2ΔSG expressing tumors (Figure 4D and 4E). Moreover, tumor metastasis could be observed with higher frequency in lateral (Left Axillary LN) but also contra-lateral (Right Axillary LN) lymph nodes for HSulf-2 expressing tumors (Figures S8C). The CS/DS chain borne by HSulf-2 is thus functionally relevant *in vivo*, at least in the context of cancer, where it attenuated the effect of the enzyme on tumor growth and metastatic invasion. In contrast, forms of HSulf-2 lacking the CS/DS chain stimulate the metastatic properties of cancer cells, thus highlighting the importance of HSulf-2 GAG modification status for considering the enzyme as a potential therapeutic target for treating human cancer.

**Fig 4:**
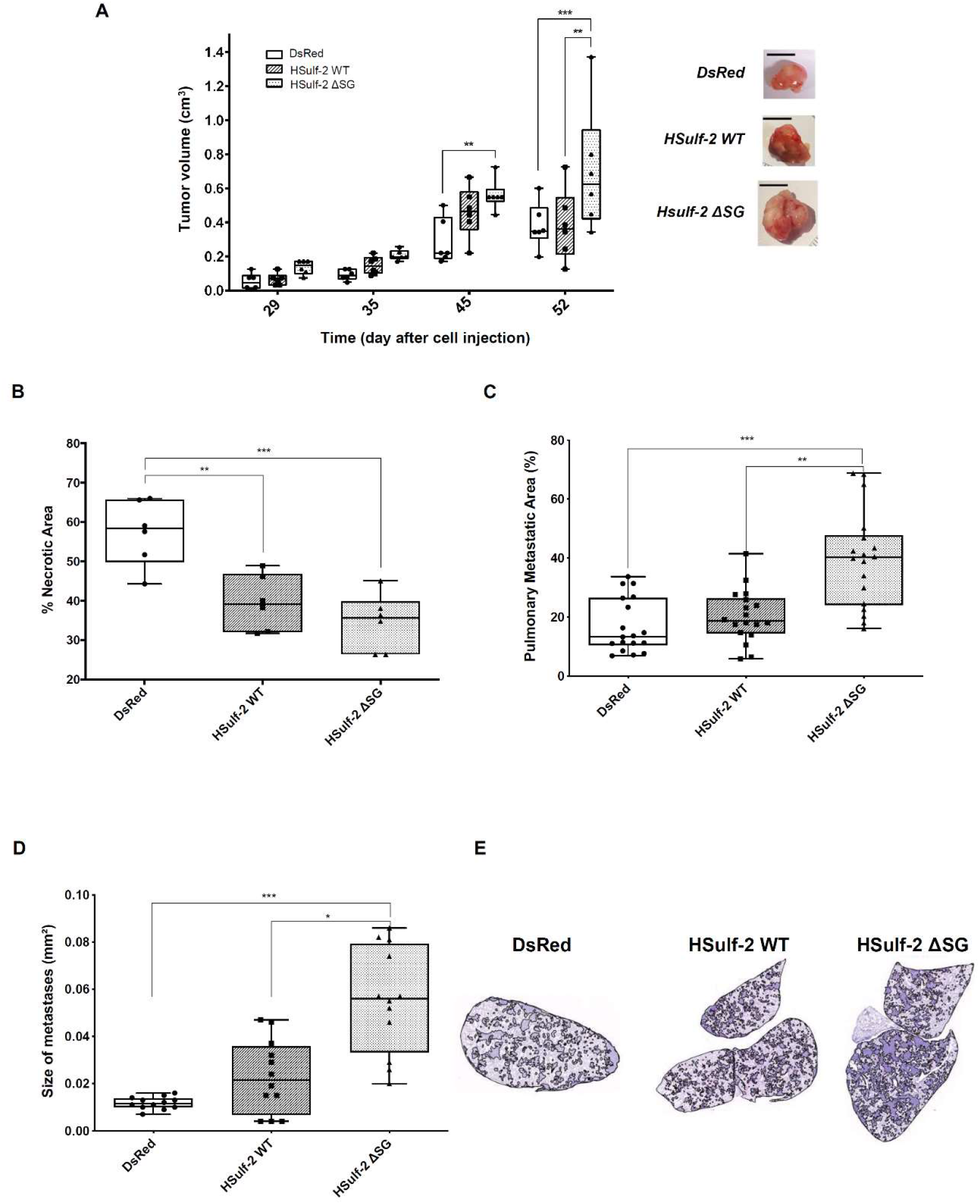
Effects of HSulf-2 WT and HSulf-2ΔSG during tumor progression and metastasis. (**A**) Time course measurement of tumor size induced by MDA-MB-231 cells expressing DsRed, HSulf-2 WT or HSulf-2ΔSG. Statistical analysis was performed using a two-way ANOVA test, ***P≤0.001 and **P<0.01. Pictures representative of each tumor group, at day 52, (A, right panel). (**B**) Histological analysis of necrotic area using eosin/hematoxin staining of tumors expressing mock DsRed, HSulf-2 WT and HSulf-2ΔSG. The percentage of necrotic area was determined on three sections from each of the six mice in each group (one-way ANOVA test, multiple comparison (Tukey’s test, n=6), ***P≤0.001 and **P<0.01). (**C**) Histological analysis of the percentage of pulmonary metastatic area from DsRed, HSulf-2 and HSulf-2ΔSG expressing tumors. The measurement was performed on three sections from each of the six mice in each group (one-way ANOVA test, multiple comparison, Tukey’s test, n=18 **P<0.01 and ***P<0.001). (**D**) The size of pulmonary metastasis in each group was quantified and analyzed as in C (n=12 **P<0.05 and ***P<0.001). (**E**) Representative images of hematoxylin/eosin stained sections of indicated lung.

## Discussion

In this study, we have shown the presence of a covalently-linked CS/DS chain on extracellular sulfatase HSulf-2, and demonstrated its functional relevance. Although CS/DS chains have been previously identified on the mucin-like domain of ADAMTS (Mead et al., 2018), the identification of HSulf-2 as a new secreted PG is unprecedented. It is well established that GAG chains provide most of PG’s biological activities, usually through the ability of the polysaccharide to bind and modulate a wide array of structural and signaling proteins. However, we show here that the GAG chain present on Hsulf-2 directly modulates its enzyme activity. These findings open new and unexpected perspectives in the understanding of the enzyme biological functions, and should contribute to clarify discrepancies in the literature.

Here, we first demonstrated that the CS/DS chain modulates HSulf-2 6-*O*-endosulfatase activity *in vitro*, most likely by competing with sulfated substrates for HS binding site occupancy, and/or through electrostatic hindrance. In support to this, we located this GAG chain on the HD domain of HSulf-2, which is critical for substrate binding. However, in an *in vivo* biological context, we anticipate that Hsulf-2 GAG chain could also modulate the enzyme function through other mechanisms. GAGs bind a wide array of cell-surface and extracellular matrix proteins. The HSulf-2 CS/DS chain could therefore promote the recruitment of GAG-binding proteins, with potentially significant functional consequences. These interactions may involve HSulf-2 in the regulation of matricrin signaling activities, or influence the diffusion and distribution of the enzyme within tissues. In line with this, our FACS-based cell-binding assay suggested enhanced attachment to the cell surface of the HSulf-2ΔSG variant *vs* the HSulf-2 WT form. Consequently, *in vivo* HSulf-2 “GAGosylation” status may not only influence the extent of HS 6-*O*-desulfation, but also the range of the enzyme activity and access to specific HS subsets in tissues.

We thus next analyzed the effect of HSulf-2 GAG chain in an *in vivo* mouse xenograft model of cancer Our data showed that overexpression of the HSulf-2ΔSG variant with enhanced activity *in vitro* promoted significantly tumor growth, vascularization and metastasis *in vivo*. The development of metastasis is a multistep process, which is a major factor of poor prognosis in cancer. Although the biological mechanisms that drive metastasis are relatively unknown, the role of cancer cell-derived matrisome proteins as prometastatic has been recently highlighted (Tian et al., 2020). Based on our data, we speculate that HSulf-2 may also participate to the extracellular cellular matrix remodeling process, and could provide an additional target to act on metastasis development. Furthermore, HSulf-2 “GAGosylation” status serve as a new metastatic promoting marker.

Beyond the field of cancer, this concept of “GAGosylation” status should prove to be critical for studying the biological functions of the Sulfs, as this may confer to the enzyme a tremendous level of functional and structural heterogeneity. It is first well known that the structure and binding properties of GAGs vary according to the biological context. We therefore anticipate further regulation of HSulf-2 catalytic activity and/or diffusion properties, depending on structural features of its CS/DS chain. In addition, our data highlighted differences in HSulf-2 furin processing amongst analyzed cell types. This could be functionally relevant, as furin maturation may affect HSulf-2 cell surface/extracellular localization as well as *in vivo* activity (Tang and Rosen, 2009). As GAGs have been previously shown to promote furin activity (Pasquato et al., 2007), we could thus hypothesize that the presence of HSulf-2 newly identified CS/DS chain at the vicinity of the two major furin cleavage sites may also influence HSulf-2 maturation status.

Here, we used a mutagenesis generated GAG-lacking HSulf-2 variant in our functional assays. However, our data suggest the co-existence of both GAG-conjugated and GAG-free HSulf-2, as PAGE analysis of GAG conjugated HSulf-2 fragments from MCF7 and HUVECs yielded distinctly different band patterns (Figure 2). The balance of expression between these two forms may therefore be critical for the control of HS 6-*O*-desulfation process. The underlying mechanisms are likely to be complex and multifactorial. Interestingly, we showed that hyaluronidases could efficiently digest Hsulf-2 GAG chain and enhance its endosulfatase activity (Figure 3E and F). Mammalian hyaluronidases are a family of 6 enzymes that catalyze the degradation of hyaluronic acid (HA) and also exhibit the ability to depolymerize CS (Csoka et al., 2001; Jedrzejas and Stern, 2005; Kaneiwa et al., 2010). Hyaluronidase expression is increased in some cancers (McAtee et al., 2014), with suggested roles in tumor invasion and tumor-associated inflammation (Dominguez Gutierrez et al., 2020; McAtee et al., 2014). However, their precise contribution remains poorly understood and contradictory. Here, we propose a new function for these enzymes, which may provide an activating mechanism of HSulf-2, through their ability to “unleash” the enzyme from its GAG chain. This perspective urges to investigate in detail the interplay of HSulf-2 and hyaluronidases during tumor progression as well as in other physiopathological conditions. In line with this, we have analyzed in details the molecular features of Hsulf-2 GAG chain and the effects of hyaluronidase on its structure and activity (Seffouh *et al*., manuscript in preparation).

Last but not least, analysis of the other human isoform HSulf-1 showed an absence of any GAG chain, at least in our HEK293 overexpressing system (Figure S3G). “GAGosylation” status could thus account for the functional differences reported between these two secreted endosulfatases.

In conclusion, we report here a most unexpected PTM of HSulf-2, by identifying the presence of a CS/DS chain on the enzyme. Our data highlight this GAG chain as a novel non-catalytic regulatory element of HSulf-2 activity, and pave the way to new directions in the study of this highly intriguing enzyme and complex regulatory mechanism of HS activity. Finally, it is worth noting that such a structurally and functionally relevant feature as a GAG chain on HSulf-2 has remained overlooked for more than 20 years. Beyond the field of the Sulfs, our findings therefore strongly encourage reconsidering afresh the importance of PTMs in complex enzymatic systems.

## Material and Methods

### Antibodies against Hsulf-2

The epitopes of antibodies against HSful-2 are summarized in Fig. S1. Polyclonal antibody H19 was newly produced by Biotem (Apprieu, France), by immunizing rabbits with a mix of 2 peptides derived from HSulf-2 sequence (C_506_DSGDYKLSLAGRRKKLF and T_563_KRHWPGAPEDQDDKDG), located with the HD domain, on each side of the furin cleavage site (see Fig. S1). Consequently, H19 is specific of the HD domain and recognizes both HSulf-2 N- and C-terminal subunits. The 2B4 monoclonal antibody, which is specific of HSulf-2 C-terminal subunit, was purchased from R&Dsystems (Mab7087). Of note, analysis of whole lysates prepared from cultured cells or tissues with 2B4 yields a sharp ~130 kDa band. This band presumably corresponds to a form in synthesis, such as non furin-processed/GAG-unmodified HSulf-2. Meanwhile, analysis of conditioned medium with 2B4 shows multiple bands corresponding to HSulf-2 unmodified or Furin/GAG-modified C-terminal subunit. The H2.3 polyclonal antibody, which is specific to the HSulf-2 N-terminal subunit, was previously described (Uchimura et al., 2006).

### Production and purification of recombinant WT and mutant HSulf-2

The expression and purification of HSulf-2 and mutants were performed as described previously (Seffouh et al., 2019a). 2019). Briefly, FreeStyle HEK293 cells (Thermo fisher scientific) were transfected with pcDNA3.1 vector encoding for HSulf-2 cDNA flanked by TEV cleavable SNAP (20.5 kDa) and His tags at N- and C-terminus, respectively. The protein was purified from conditioned medium by cation exchange chromatography on a SP sepharose column (GE healthcare) in 50 mM Tris, 5 mM MgCl_2_, 5 mM CaCl_2_, pH 7.5, using a 0.1-1 M NaCl gradient, followed by size exclusion chromatography (Superdex200, GE healthcare) in 50 mM Tris, 300 mM NaCl, 5 mM MgCl_2_, 5 mM CaCl_2_, pH 7. Treatment of HSulf-2 with chondroitinase ABC was achieved by incubating 250 µg of enzyme with 100mU chondroitinase ABC (Sigma) overnight at 4°C. HSulf-2ΔSG mutants (ΔSG, ΔSG1, ΔSG2) were generated by site directed mutagenesis (ISBG Robiomol platform) and purified as above.

### Analysis of HSulf-2 expression

MDA-MB-231 cells were lysed with RIPA buffer for 2 h at 4°C and tissues were disrupted and lysed in RIPA buffer (Sigma-Aldrich) using a MagNA Lyser instrument (Roche) with ceramic beads. Supernatants were collected and protein concentration was determined using a BCA protein Assay kit (Thermo Scientific). Cell lysates (eq. of 3.10^5^ cells), tumor lysates (eq. of 50 µg of total proteins) or purified recombinant proteins were then separated by SDS-PAGE, followed by transfer onto PVDF membrane. Proteins were probed using either rabbit polyclonal H19 (dil. 1/1000) or mouse monoclonal 2B4 (dil. 1/500) antibodies, followed by incubation with HRP-conjugated anti-rabbit (Thermo Scientific, dil. 1/5000), anti-mouse (Thermo Scientific, dil. 1/5000) secondary antibodies.

Endogenous CS/DS modification of HSulf-2 was analyzed in two cell lines: the MCF-7 human breast cancer cells and human umbilical vein endothelial cells (HUVECs). MCF-7 cells were cultured at 37 °C for 48 h in OPTI-MEM, after which culture medium was collected and concentrated on Amicon Ultra Filters (30 kDa cut-off, Millipore, Burlington, MA). Conditioned medium from MDA-MB231 cells was prepared likewise, using FreeStyle medium instead of OPTI-MEM. HSulf-2 in concentrated samples were analyzed by Western blotting as described below. HUVECs were cultured at 37 °C in OPTI-MEM containing 0.5 % FBS for 24 h, after which culture medium was collected and concentrated on Amicon Ultra Filters. Concentrated samples were incubated with GlcNAc-binding wheat germ agglutinin (WGA)-coated beads (Vector Laboratories, Burlingame, CA) at 4 °C overnight, and proteins that were captured by WGA beads were analyzed by Western blotting. For elimination of CS/DS chains, the concentrated MCF-7 culture media or WGA bead-bound materials were treated with chondroitinase ABC (1 U/mL) or heat-inactivated chondroitinase ABC (1 U/mL), at 37 °C for 1 h. Proteins in the samples were separated by SDS-PAGE with 5–20% gels (Wako Pure Chemical Industries, Osaka, Japan) and were transferred to PVDF membranes. HSulf-2 proteins were probed with the 2B4 mouse monoclonal anti-HSulf-2 antibody (dil. 1/500) or the H2.3 rabbit polyclonal anti-HSulf-2 antibody (dil. 1/500) followed by a horseradish peroxidase-labeled anti-mouse or rabbit IgG antibody (Cell Signaling Technology, Danvers, MA) and ImmunoStar LD (Wako Pure Chemical Industries). Signals were visualized by using a LuminoGraph image analyzer (ATTO, Tokyo, Japan).

### SAXS analysis

SAXS data were collected at the European Synchrotron Radiation Facility (Grenoble, France) on the BM29 beamline at BioSAXS. The standard energy as set to 12.5 keV and a Pilatus 1M detector was used to record the scattering patterns. The sample-to-detector distance was set to 2.867m (q-range is 0.025 - 6 nm-1). Samples were set in quartz glass capillary with an automated sample changer. The scattering curve of the buffer (before and after) solution was subtracted from the sample’s SAXS curves. Scattering profiles were measured at several concentrations, from 0.5 to 1.5 mg/mL at room temperature. Data were processed using standard procedures with the ATSAS v2.8.3 suite of programs (Petoukhov et al., 2012). The *ab initio* determination of the molecular shape of the proteins was performed as previously described (Pérard et al., 2018). Radius of gyration (Rg) and forward intensity at zero angle (I(0)) were determined with the programs PRIMUS (Konarev et al., 2003), by using the Guinier approximation at low Q value, in a Q.Rg range < 1.5:

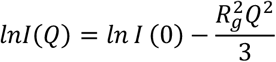

Porod volumes and Kratky plot were determined using the Guinier approximation and the PRIMUS programs. The pairwise distance distribution function P(r) were calculated by indirect Fourier transform with the program GNOM (Svergun, 1992). The maximum dimension (Dmax) value was adjusted in order that the Rg R value obtained from GNOM agreed with that obtained from Guinier analysis. In order to build *ab initio* models, several independent DAMMIF (Franke and Svergun, 2009) models were calculated in slow mode with pseudo chain option and merged using the program DAMAVER (Konarev et al., 2003).

### MALDI-TOF MS analysis of HSulf-2ΔSG

MS experiments were carried out on a MALDI Autoflex speed TOF/TOF MS instrument (Bruker Daltonics, Germany), equipped with a SmartBeam II™ laser pulsed at 1 kHz. The spectra were recorded in the positive linear mode (delay: 600 ns; ion source 1 (IS1) voltage: 19.0 kV; ion source 2 (IS2) voltage: 16.6 kV; lens voltage: 9.5 kV). MALDI data acquisition was carried out in the mass range 5000– 150000 Da, and 10000 shots were summed for each spectrum. Mass spectra were processed using FlexAnalysis software (version 3.3.80.0, Bruker Daltonics). The instrument was calibrated using mono- and multi-charged ions of BSA (BSA Calibration Standard Kit, AB SCIEX, France). HSulf-2ΔSG was desalted as previously described (Seffouh et al., 2019b). MALDI-TOF MS analysis was achieved by mixing 1.5 μL of sinapinic acid matrix at 20 mg/mL in acetonitrile/water (50/50; v/v), 0.1% TFA, with 1.5 μL of the desalted protein solution (0.57 mg/mL).

### LC-MS/MS identification of Hsulf-2 GAG chain and its attachment site

The glycoproteomics protocol used for characterizing proteoglycans has been published earlier(Noborn et al., 2015) and most recently summarized in detail for analyses of CS proteoglycans of human cerebrospinal fluid (Noborn et al Methods in Molecular Biology, in press). In the present work, the starting material was conditioned cell media, without fetal calf serum, obtained from SH-SY5Y cells kindly provided by Drs. Thomas Daugbjerg-Madsen and Katrine Schjoldager, University of Copenhagen, Denmark.

### In vitro enzyme activity assays

Detailed protocols for arylsulfatase and endosulfatase assays have been described elsewhere (Seffouh et al., 2019a). For the arylsulfatase assay, the enzyme (2 µg) was incubated for 4h with 10 mM 4MUS (Sigma) in 50 mM Tris, 10 mM MgCl_2_ pH 7.5 for 1-4 h at 37°C, and the reaction was followed by fluorescence monitoring (excitation 360 nm, emission 465 nm). Results are expressed as a fold of fluorescence increase compared to negative control (4MUS alone) and corresponds to means +/-SD of three independent experiments. The endosulfatase assay was achieved by incubating 25 µg of Heparin with 3 µg of enzymes in 50 mM Tris, 2.5 mM MgCl_2_ pH 7.5, for 4 h at 37°C. Disaccharide composition of Sulf-treated heparin was then determined by exhaustive digestion of the polysaccharide (48 hours at 37°C) with a cocktail of heparinase I, II and III (10 mU each), followed by RPIP-HPLC analysis (Henriet et al., 2017), using NaCl Gradients calibrated with authentic standards (Iduron). Hsulf-2 (2 µg) digestion with hyaluronidase (Sigma/Aldrich, 2µg) was achieved after a 2 h incubation in 50 mM Tris pH 7.5 at 37°C, prior to the endosulfatase assay, which was performed as above. Incubation of heparin with hyaluronidase alone showed no effect on the disaccharide analysis (data not shown).

### Analysis of HSulf-2/heparin binding immuno-assay

As reported before (Seffouh et al., 2019a), microliter plates were first coated with 10 µg/ml streptavidin in TBS buffer, then incubated with 10 µg/ml biotinylated heparin, and saturated with 2% BSA. All the incubations were achieved for 1h at RT, in 50 mM Tris-Cl, 150 mM NaCl, pH 7.5 (TBS) buffer. Next, the recombinant protein was added, then probed with H2.3 primary rabbit polyclonal anti-HSulf-2 antibody (dil. 1/1000) followed by fluorescent-conjugated secondary anti-rabbit antibody (Jackson ImmunoResearch Laboratories, dil. 1/500). All the incubations were performed for 2 h at 4 °C in TBS, 0.05% Tween, and were separated by extensive washes with TBS, 0.05% Tween. Finally, fluorescence of each well was measured (excitation 485 nm, emission 535 nm). K_D_s were determined by Scatchard analysis of the binding data. Results shown are representative of three independent experiments.

### FACS analysis

Wish cells (1million for each condition) were washed with PBS, 1% BSA (the same buffer is used all along the experiment), and incubated with 5 µg of HSulf-2 enzymes (2 h at 4°C). After extensive washing, cells were incubated with H2.3 primary antibody (dil. 1/500, 1 h at 4°C), washed again, then with secondary AlexA 488-conjugated antibody (Jackson ImmunoResearch Laboratories, dil. 1/500, 1 h at 4°C). FACS analysis of cell fluorescence was performed on a MACSQuant Analyzer (Miltenyi Biotec, excitation 485 nm, emission 535 nm) by calculating median over 25000 events. Data are represented as means +/-SD of three independent experiments.

### Lentiviral transduction of MDA-MB 231 cells

HSulf-2 (WT and variant) encoding cDNAs were cloned into the pLVX lentiviral vector (Clonetech). This vector was then used in combination with viral vectors GAG POL (psPAX2) and ENV VSV-G (pCMV) to transduce HEK293T and produce lentiviruses released in the extracellular medium. The pLVX-Ds-Red N1 (Clonetech) was used as negative control.

MDA-MB-231 cells were purchased from ATCC and were cultured in Leibovitz’s medium (Life Technologies) supplemented with 10% fetal bovine serum, 100 U/ml of penicillin, 100 µg/ml of streptomycin (Life Technologies). For infection, MDA-MB-231 cells were plated into 6 well-plates (8 x 10^5^ in 2 mL of serum-supplemented Leibovitz’s medium). The day after, adherent cells were incubated with 1 ml of lentiviral medium diluted in 1 mL of serum-supplemented medium containing 8 μg/μL of polybrene (Sigma/Aldrich). After 4 h, 1 mL of medium were added to cultures and transduction was maintained for 16h before washing the cells and changing the medium. For stable transduction, puromycin selection was started 36 h post-infection (at the concentration of 2 μg/mL, Life Technologies) and was maintained thereafter.

### In vivo experiments

*In vivo* experiment protocols were approved by the institutional guidelines and the European Community for the Use of Experimental Animals. 7-weeks-old female NOD SCID GAMMA/J mice were purchased from Charles River and maintained in the Animal Resources Centre of our department. 1×10^6^ MDA-MB-231 cells resuspended in 50 % MatrigelTM (Becton Dickinson) in Leibovitz medium (Life Technologies) were injected into the fat pad of #4 left mammary gland. Tumor growth was recorded by sequential determination of tumor volume using caliper. Tumor volume was calculated according to the formula V = ab^2^/2 (a, length; b, width). Mice were euthanized after 52 days through cervical dislocation. Tumors and axillary lymph nodes were collected, weighed and either fixed for 2h in 4 % paraformaldehyde (PFA) and embedded in paraffin, or stored at −80°C for WB analysis. Tissue necrosis was assessed by Hematoxylin/eosin staining and ImageJ quantification. For vascularization analysis, sections (5μm thick) of formalin-fixed, paraffin embedded tumor tissue samples were dewaxed, rehydrated and subjected to antigen retrieval in citrate buffer (Antigen Unmasking Solution, Vector Laboratories) with heat. Slides were incubated for 10 min in hydrogen peroxide H_2_O_2_ to block endogenous peroxidases and then 30 min in saturation solution (Histostain, Invitrogen) to block non-specific antibody binding. This was followed by overnight incubation, at 4°C, with primary antibody against αSMA (Ab124964, Abcam, dil. 1/500). After washing, sections were incubated with a suitable biotinylated secondary antibody (Histostain kit, Invitrogen) for 10 min. Antigen-antibody complexes were visualized by applying a streptavidin-biotin complex (Histostain, Invitrogen) for 10 min followed by NovaRED substrate (Vector Laboratories). Sections were counterstained with hematoxylin to visualize nucleus. Control sections were incubated with secondary antibody alone. Lungs were inflated using 4% PFA and embedded in paraffin. The metastatic burden was assessed by serial sectioning of the entire lungs, every 200µm. Hematoxylin and eosin staining was performed on lung and lymph nodes sections (5 µm thick). Images were acquired using AxioScan Z1 (Zeiss) slide scanner and quantified using FiJi software.

### Statistical analysis

Experimental data are shown as mean ± standard error of the mean (SEM) unless specified otherwise. Comparisons between multiple groups were carried out by a repeated-measures two-way analysis of variance (ANOVA) with Tukey’s multiple comparisons test to evaluate the significance of differential tumor growth between three groups of mice; an ordinary two-way ANOVA with Bonferroni’s test and an ordinary one-way ANOVA were carried out to evaluate *in vitro* activity and binding of HSulf-2, the differential level of necrosis and vascularization inside tumors, and pulmonary metastases (number and area). Prism 6 (GraphPad Software, Inc., CA) was used for analyses. Probability value of less than 0.05 was considered to be significant. * P < 0.05, ** P < 0.01, *** P < 0.001 and **** P < 0.0001.

## Acknowledgments

The authors would like to thank the animal unit staff (Jeannin I., Bama S., Magallon C., Chaumontel N. and Pointu H.) at the Interdiciplinary Research Institute of Grenoble (IRIG) for animal husbandry. This work used the platforms of the Grenoble Instruct-ERIC center (ISBG; UMS 3518 CNRS-CEA-UJF-EMBL) within the Grenoble Partnership for Structural Biology (PSB). Platform access was supported by FRISBI (ANR-10-INBS-05-02) and GRAL, a project of the University Grenoble Alpes graduate school (Ecoles Universitaires de Recherche) CBH-EUR-GS (ANR-17-EURE-0003). This work was also supported by the CNRS and the GDR GAG (GDR 3739), the “Investissements d’avenir” program Glyco@Alps (ANR-15-IDEX-02), by grants from the Agence Nationale de la Recherche (ANR-12-BSV8-0023 and ANR-17-CE11-0040) and Université Grenoble-Alpes (UGA AGIR program), the Swedish Research Council (2017-00955 to GL and to the Swedish National Infrastructure for Biological Mass Spectrometry (BIOMS)), and the Inga-Britt and Arne Lundbergs Forskningsstiftelse. IBS acknowledges integration into the Interdisciplinary Research Institute of Grenoble (IRIG, CEA).

## Supplementary material

**Fig S1:**
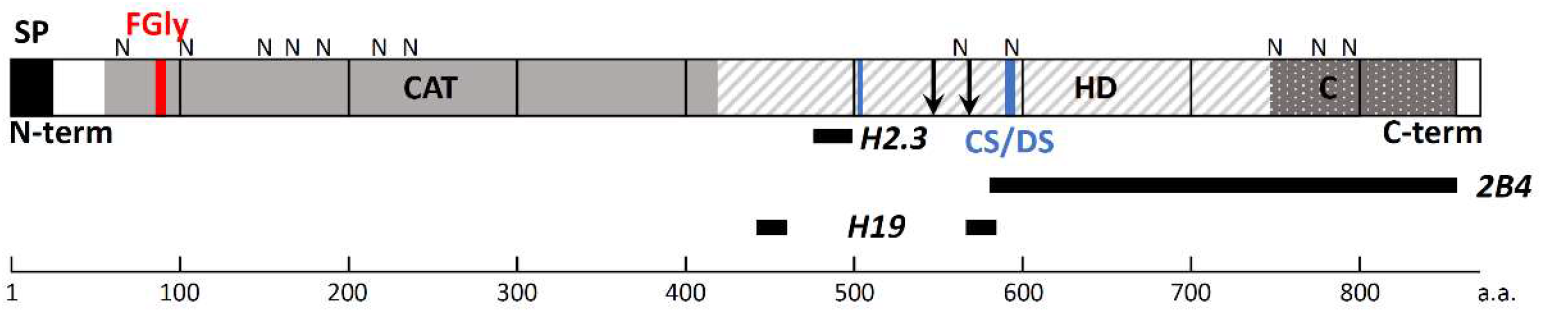
Schematic representation of HSulf-2 molecular organization, PMTs and antibody epitopes. HSulf-2 870 amino-acid (a.a.) pro-protein comprises a signal peptide (SP, black box) and a polypeptide processed through Furin cleavage (black arrows) into two N-terminal (N-term) and C-terminal (C-term) subunits. HSulf-2 comprises two major functional domains: a catalytic domain (CAT, in grey) and a highly basic hydrophilic region (HD, hatched in grey), and features a C-terminal region sharing homology with glucosamine-6-sulfatase homolog (C, dotted). Potential *N*-glycosylation sites (N), the catalytic FGly residue (FGly, in red and bold) and the SG dipeptides (blue, in bold for S_583_G) and antibody epitopes (black bars) are indicated.

**Fig S2:**
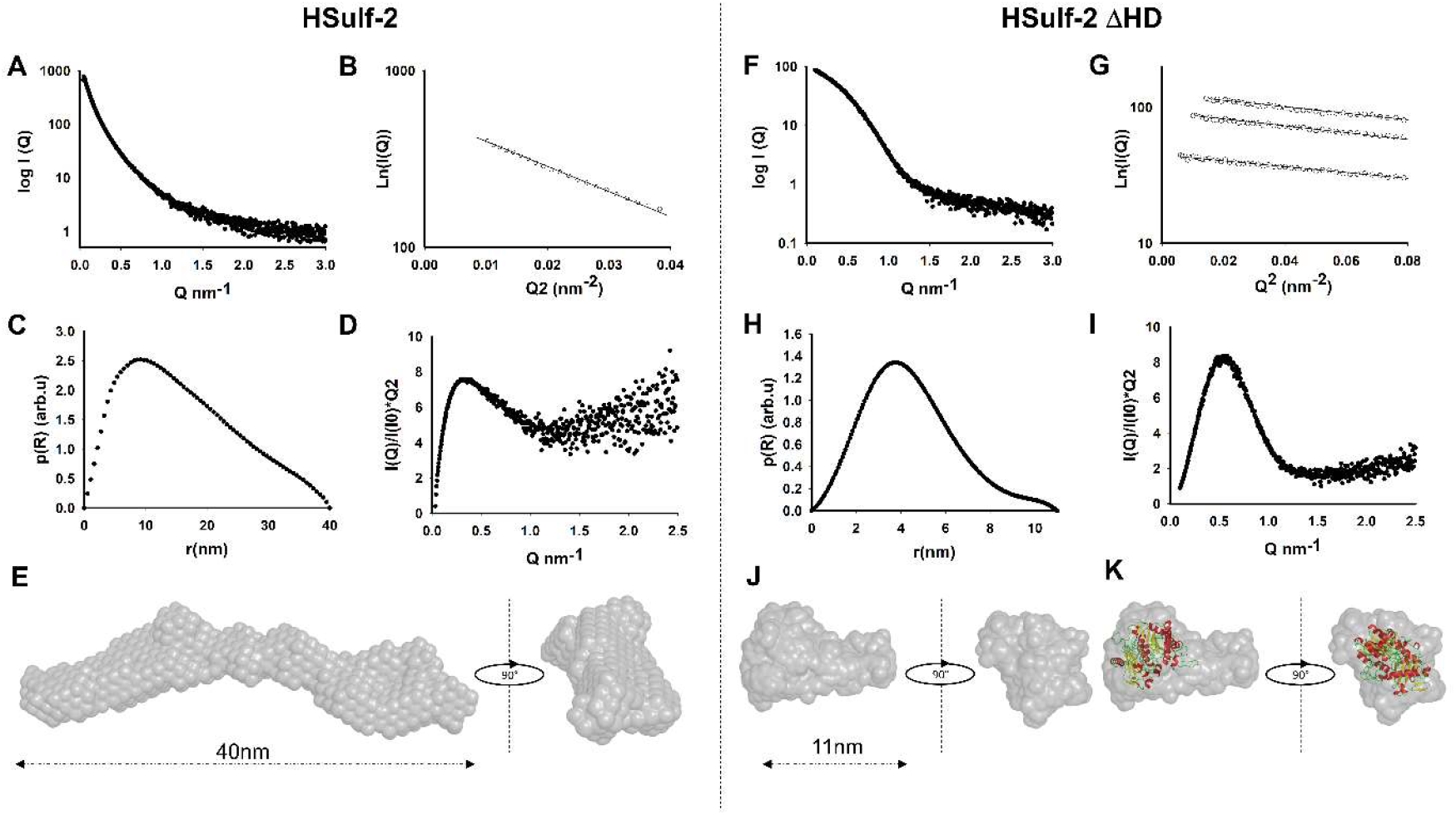
Study of HSulf-2 and HSulf-2 ΔHD by Small Angle X-Ray scattering. SAX Analysis of HSulf-2 (Panels **A-E**) and HSulf-2 ΔHD (Panels **F-K**). (**A**) Scattering curves of experimental data of HSulf-2 in solution. (**B**) Linear dependence of ln[I(Q)] vs Q2 determined by Guinier plot at 0.6mg/ml with a Rg of 13nm and I0 of 700 with a porod volume more than 1000. HSulf-2 give a MW_exp_: 700 kDa. MW_malls_: 1000 kDa, MW_th protein_: 98,53 kDa. This data indicate the presence of elongated molecule with a potential rode shape. (**C**) Pair distribution function p(R) in arbitrary units (arb.u) *vs. r* (nm) determined by GNOM with a D_max_ of 40 nm +/-3 nm indicate that the HSulf-2 is an elongated molecule in solution. (**D**) I globularity and flexibility analysis of HSulf-2. Kratky plot(I(q)*q^2^ vs. q) of HSulf-2 not converge to the q axis witch and indicate the presence of mixture of multidomain protein with flexible linker and unfolded region (could be allocate to the GAG). (**E**) Final *ab initio* model of HSulf-2 generated with individual DAMMIF model in slow mode. DAMAVER classification under NSD value indicates the presence of several clusters (NSD > 1.5) for HSulf-2 suggesting the presence of flexible regions. The proposed final model of HSulf-2 combines 14 of the 49 models calculated with the best NSD (between 1.5 to 1.7). (**F**) Scattering curves of experimental data of HSulf-2 ΔHD domain in solution. (**G**) Linear dependence of ln[I(Q)] vs Q^2^ determined by Guinier plot at several concentrations between 0,5 to 3mg/ml give a linear region with Rg of 3.9nm and a I0 of 87 with a porod volume of 144. MW_exp_: 87 kDa (MW_malls_: 84 kDa, MW_th protein_: 64 kDa). This data indicate the presence of potential globular protein. (**H**) Pair distribution function p(R) in arbitrary units (arb.u) *vs. r* (nm) determined by GNOM give a D_max_ of 11 nm with a relative globular shape. (**I**) Kratky plot of HSulf-2 ΔHD present a “bell-shape” peak at low q and converges to the q axis at high q corresponding to a well-folded globular protein. (**J**) Final *ab initio* model of HSulf-2 ΔHD generated with individual DAMMIF model in slow mode and merged with Damaver (NSD < 0.7). The HSulf-2 ΔHD *ab initio* model give a globular envelope with a small-elongated part. (**K**) Superimposition of prediction structure of HSulf-2 ΔHD based on pdb: 4UPL (from Silicibacter pomeroyi) structure (PHYRE analysis) into HSulf-2 ΔHD SAXS envelope with Supcomb20 program.

**Fig S3:**
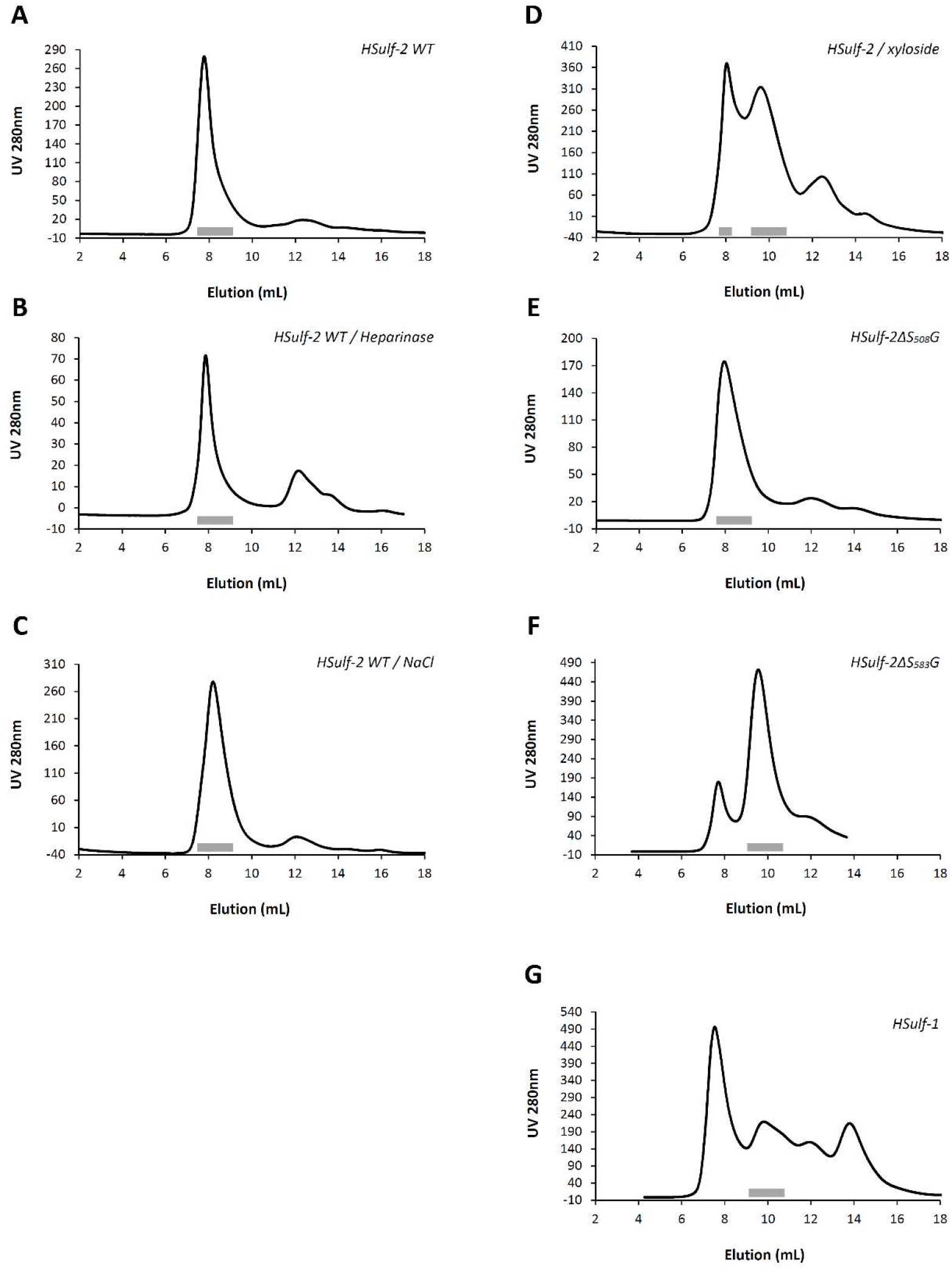
Size-exclusion chromatography of HSulf-2 WT, HSulf-2 variants and HSulf-1. Size exclusion chromatography profile of HSulf-2 WT without (**A**), or following pre-treatment with heparinase I, II, III (**B**), 2 M NaCl (**C**), or xyloside (**D**). Size exclusion chromatography profile of HSulf-2 ΔS_508_G (**E**), ΔS_583_G (**F**), or HSulf-1 (**G**). Grey bars show Sulf-containing peaks.

**Fig S4:**
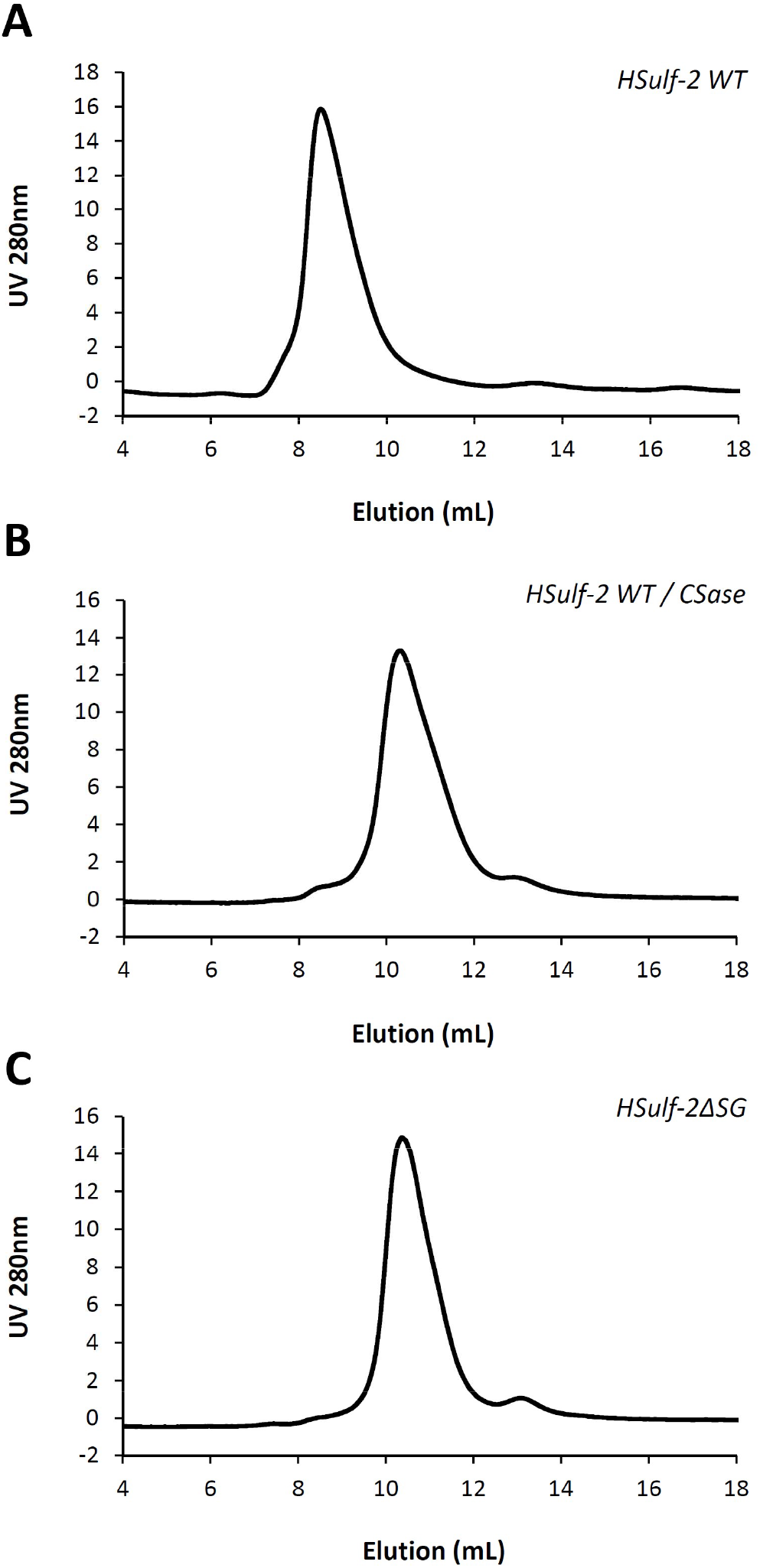
Size-exclusion chromatography of TEV-treated HSulf-2 WT and variant. Size exclusion chromatography profile of TEV-treated HSulf-2 WT (**A**), chondroitinase ABC-treated HSulf-2 WT (**B**), or HSulf-2ΔSG (**C**).

**Fig S5:**
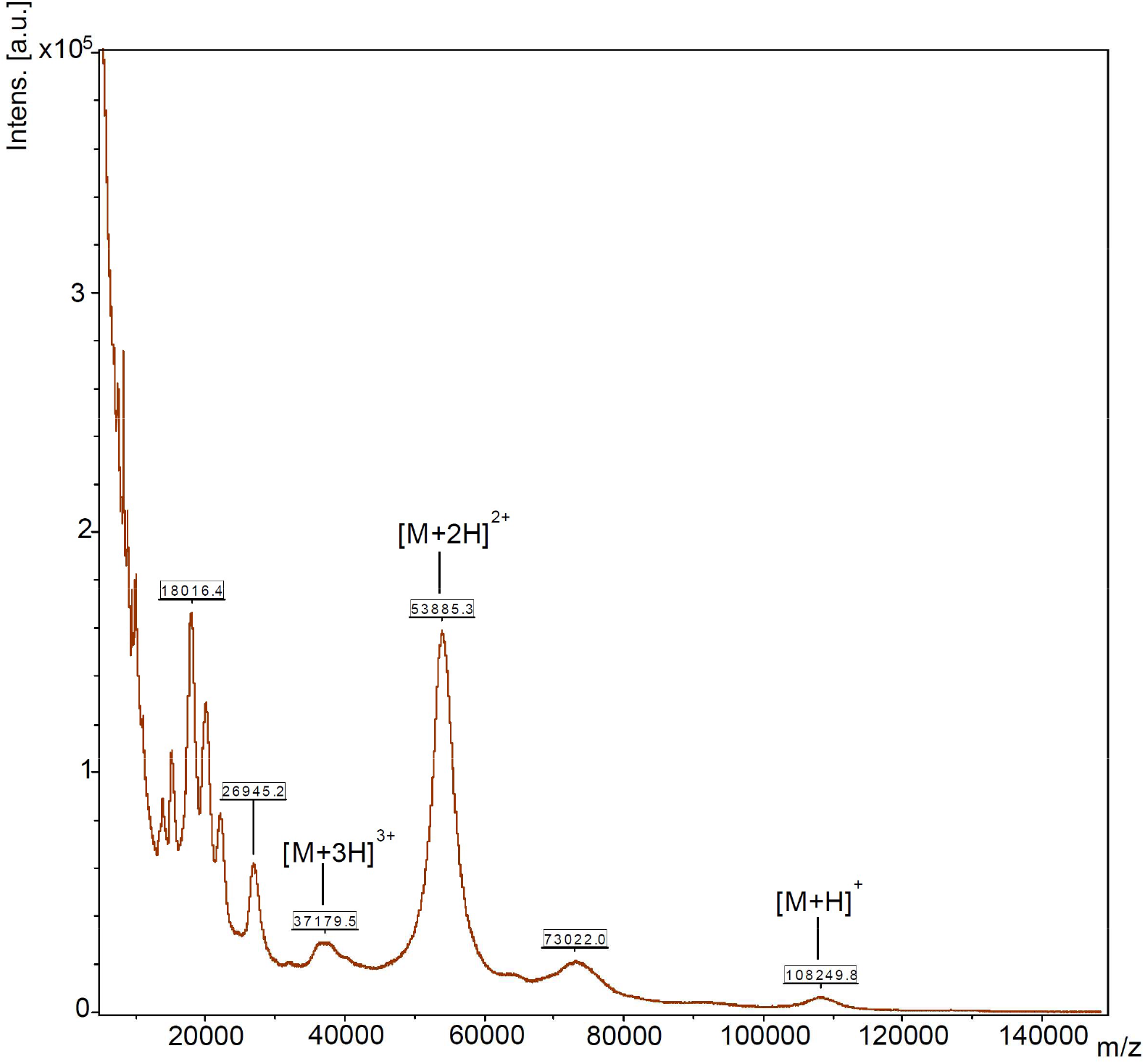
MALDI-TOF mass spectrometry analysis of HSulf-2ΔSG. Mass spectrum of HSulf-2ΔSG in positive ionization mode (100 kDa-filtrated HSulf-2ΔSG mixed with sinapinic acid matrix, linear mode). HSulf-2ΔSG is detected as the protonated species [M+H]^+^ and corresponding doubly and tri-charged [M+2H]^2+^ and [M+3H]^3+^ ions.

**Fig S6:**
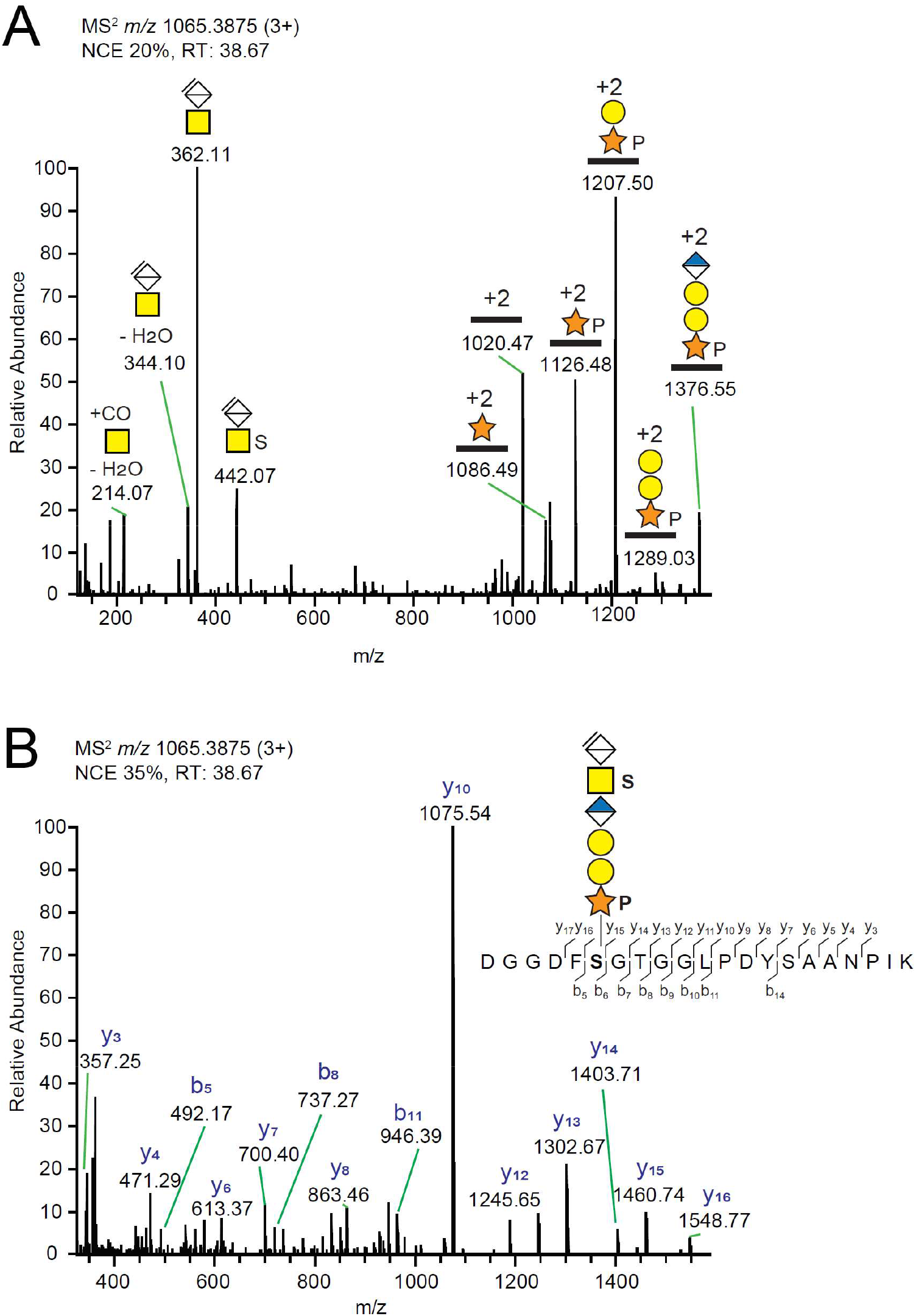
LC-MS/MS detection of a CS/DS GAG linkage region attached to S_583_ of HSulf-2. HSulf-2 glycopeptides were obtained by trypsin digestion of media of cultured SH-SY5Y cells, followed by enrichment on a SAX column, and thereafter treatment with chondroitinase ABC. The spectral files were filtered for the MS^2^ diagnostic ion at *m/z* 362.1083 corresponding to the delta-hexuronic acid - *N*-acetylgalactosamine disaccharide, common to all CS/DS linkage region glycopeptides. (**A**) MS^2^ spectrum of the 578-DGGDFSGTGGLPDYSAANPIK-598 glycopeptide obtained by HCD with normalized collision energy of 20%, providing prominent glycan fragmentations. (**B**) MS^2^ spectrum of the same glycopeptide obtained at normalized collision energy of 35%, displaying peptide sequence fragmentation with b- and y-ions annotated in the sequence. The positioning and distinction of sulfate (79.9568 u) and phosphate (79.9663 u) modifications were done by manually interpreting the MS^2^ spectra. The MS^2^ spectrum thus displayed a mass shift of 79.9570 u between *m/z* 362.1083 and *m/z* 442.0653, demonstrating the presence of a sulfate modification on the GalNAc residue. A mass shift of 212.0084 u was observed between *m/z* 1019.9724 (2+) and m/z 1125.9766 (2+), demonstrating the presence of a xylose plus phosphate modification of the peptide (the theoretical mass of this modification is 212.0086 u (132.0423 u + 79.9663 u).

**Fig S7:**
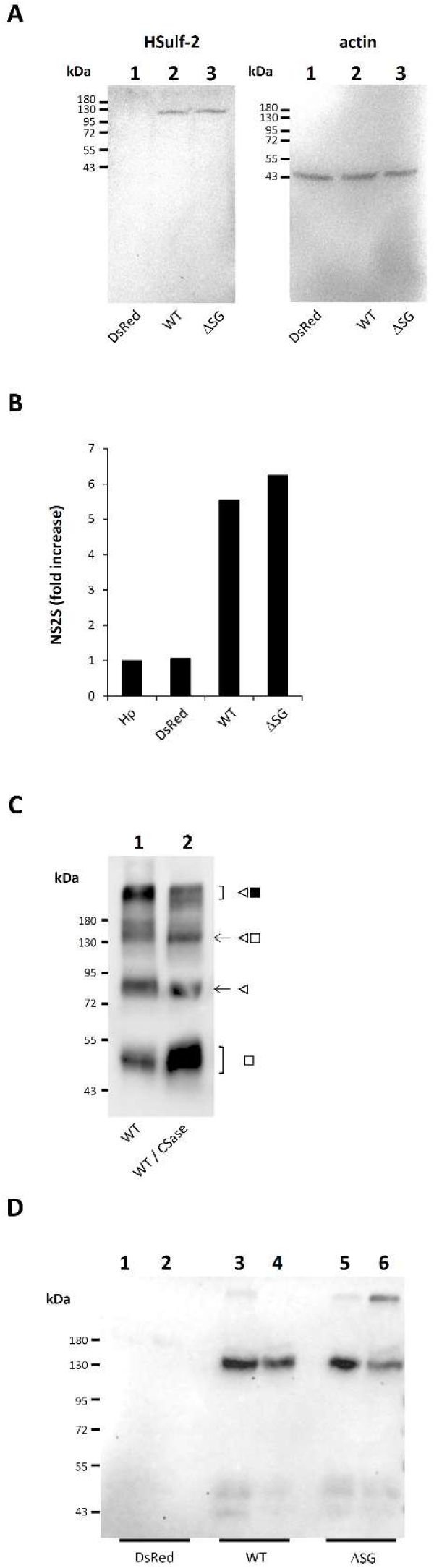
Expression and activity of HSulf-2 in MDA-MB-231 transduced cells. (**A**) Western blot analysis of Dsred (lanes 1), HSulf-2 WT (lanes 2) or HSulf-2ΔSG (lanes 3) transduced MDA-MB-231 cell lysates, using anti C-terminal HSulf-2 2B4 antibody and anti-actin antibody (Sigma, ref A-2066) as a loading control. (**B**) Endosulfatase activity was monitored by treating heparin with Dsred, HSulf-2 WT or HSulf-2ΔSG transduced MDA-MB-231 cell conditioned medium. Results are expressed as fold increase of NS2S disaccharide content compared to untreated heparin (control). (**C**) Western blot analysis (H19 antibody) of HSulf-2 WT transduced MDA-MB-231 cell conditioned medium prior to (WT, lane 1) or after (WT / CSase, lane 2) treatment with chondroitinase ABC. (**D**) Western blot analysis (2B4 antibody) of mice tumor lysates resulting from injections of DsRed (lanes 1 and 2), HSulf-2 WT (lanes 3 and 4) or HSulf-2ΔSG (lanes 5 and 6) transduced MDA-MB-231 cells.

**Fig S8:**
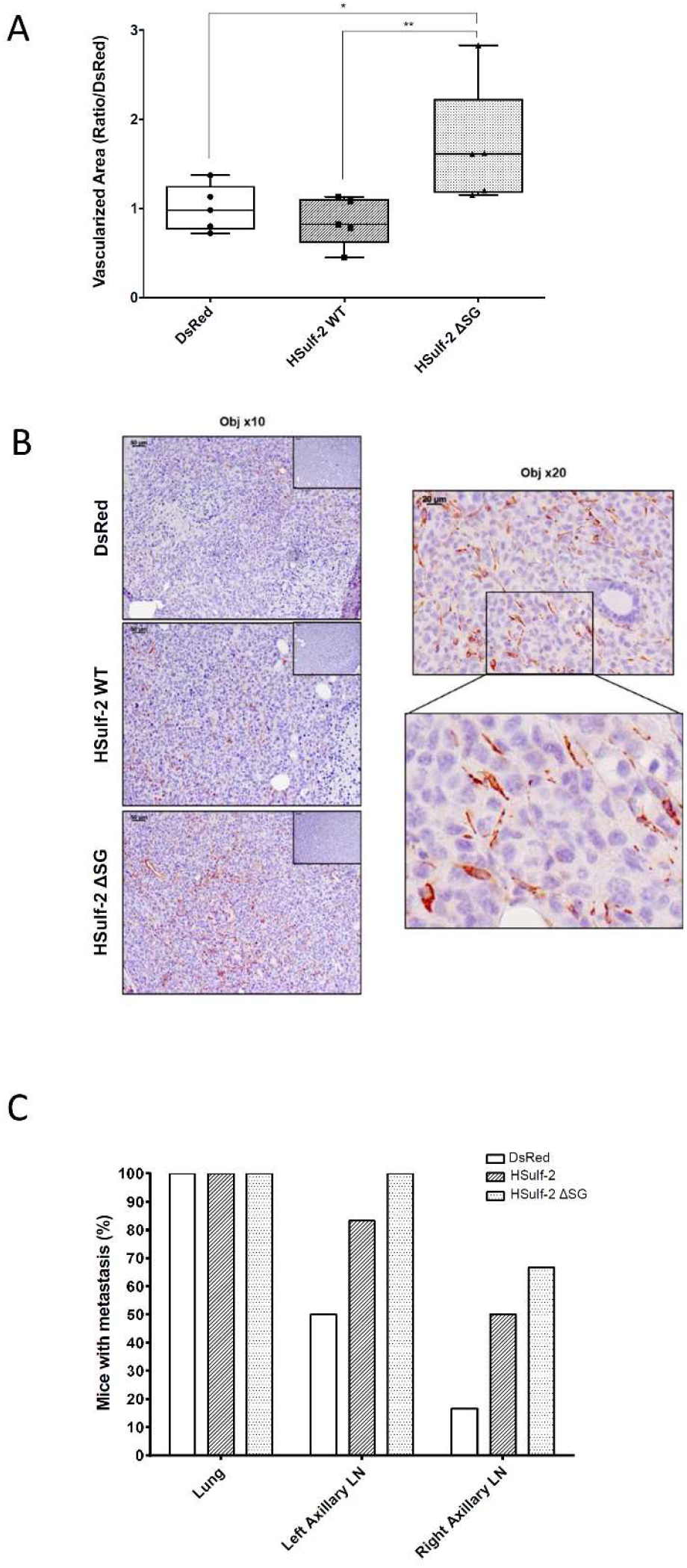
Effects of HSulf-2 WT and HSulf-2ΔSG on tumor metastasis. (**A**) Histological analysis of the vascularized area, using α Smooth Muscle Actin (αSMA) immunostaining of tumors expressing mock DsRed, HSulf-2 WT and HSulf2ΔSG. The calculation of vascularized area of tumors was performed on five mice in each group. For each mouse, 4 ROIs (Region Of Interest) were quantified: The αSMA positive area was measured for each ROI and divided by the total ROI’s area giving for each mice a percentage of vascularized area. In each group, the median of this percentage was divided by the mean of the median of the 5 DsRed control mice (one-way ANOVA, multiple comparison, Bonferroni test,*P=0.07, **P=0.03). (**B**) Representative images of αSMA staining with different magnifications. (**C**) Percentage of mice with metastasis found in the lung, left axillary and right axillary lymph node (LN) from DsRed, HSulf-2 and HSulf-2ΔSG expressing tumors.

## References

Ai, X., Do, A.T., Kusche-Gullberg, M., Lindahl, U., Lu, K., and Emerson, C.P., Jr. (2006). Substrate specificity and domain functions of extracellular heparan sulfate 6-O-endosulfatases, QSulf1 and QSulf2. J Biol Chem 281, 4969–4976.

Ambasta, R.K., Ai, X., and Emerson, C.P., Jr. (2007). Quail Sulf1 function requires asparagine-linked glycosylation. J Biol Chem 282, 34492–34499.

Bret, C., Moreaux, J., Schved, J.F., Hose, D., and Klein, B. (2011). SULFs in human neoplasia: implication as progression and prognosis factors. J Transl Med 9, 72.

Chua, J.S., and Kuberan, B. (2017). Synthetic Xylosides: Probing the Glycosaminoglycan Biosynthetic Machinery for Biomedical Applications. Acc. Chem. Res. 50, 2693–2705.

Csoka, A.B., Frost, G.I., and Stern, R. (2001). The six hyaluronidase-like genes in the human and mouse genomes. Matrix Biol. 20, 499–508.

Dhoot, G.K., Gustafsson, M.K., Ai, X., Sun, W., Standiford, D.M., and Emerson, C.P., Jr. (2001). Regulation of Wnt signaling and embryo patterning by an extracellular sulfatase. Science 293, 1663–1666.

Dierks, T., Dickmanns, A., Preusser-Kunze, A., Schmidt, B., Mariappan, M., von Figura, K., Ficner, R., and Rudolph, M.G. (2005). Molecular basis for multiple sulfatase deficiency and mechanism for formylglycine generation of the human formylglycine-generating enzyme. Cell 121, 541–552.

Dominguez Gutierrez, P.R., Kwenda, E.P., Donelan, W., O’Malley, P., Crispen, P.L., and Kusmartsev, S. (2020). Hyal2 expression in tumor-associated myeloid cells mediates cancer-related inflammation in bladder cancer. Cancer Res.

El Masri, R., Seffouh, A., Lortat-Jacob, H., and Vivès, R.R. (2017). The “in and out” of glucosamine 6-O-sulfation: the 6th sense of heparan sulfate. Glycoconj. J. 34, 285–298.

Esko, J.D., and Zhang, L. (1996). Influence of core protein sequence on glycosaminoglycan assembly. Curr. Opin. Struct. Biol. 6, 663–670.

Franke, D., and Svergun, D.I. (2009). DAMMIF, a program for rapid ab-initio shape determination in small-angle scattering. J. Appl. Crystallogr. 42, 342–346.

Frese, M.A., Milz, F., Dick, M., Lamanna, W.C., and Dierks, T. (2009). Characterization of the human sulfatase Sulf1 and its high affinity heparin/heparan sulfate interaction domain. J Biol Chem 284, 28033–28044.

Hanson, S.R., Best, M.D., and Wong, C.H. (2004). Sulfatases: structure, mechanism, biological activity, inhibition, and synthetic utility. Angew Chem Int Ed Engl 43, 5736–5763.

Harder, A., Möller, A.-K., Milz, F., Neuhaus, P., Walhorn, V., Dierks, T., and Anselmetti, D. (2015). Catch bond interaction between cell-surface sulfatase Sulf1 and glycosaminoglycans. Biophys. J. 108, 1709–1717.

Henriet, E., Jäger, S., Tran, C., Bastien, P., Michelet, J.-F., Minondo, A.-M., Formanek, F., Dalko-Csiba, M., Lortat-Jacob, H., Breton, L., et al. (2017). A jasmonic acid derivative improves skin healing and induces changes in proteoglycan expression and glycosaminoglycan structure. Biochim. Biophys. Acta 1861, 2250–2260.

Jedrzejas, M.J., and Stern, R. (2005). Structures of vertebrate hyaluronidases and their unique enzymatic mechanism of hydrolysis. Proteins Struct. Funct. Bioinforma. 61, 227–238.

Kaneiwa, T., Mizumoto, S., Sugahara, K., and Yamada, S. (2010). Identification of human hyaluronidase-4 as a novel chondroitin sulfate hydrolase that preferentially cleaves the galactosaminidic linkage in the trisulfated tetrasaccharide sequence. Glycobiology 20, 300–309.

Konarev, P.V., Volkov, V.V., Sokolova, A.V., Koch, M.H.J., and Svergun, D.I. (2003). PRIMUS: a Windows PC-based system for small-angle scattering data analysis. J. Appl. Crystallogr. 36, 1277–1282.

Lemjabbar-Alaoui, H., van Zante, A., Singer, M.S., Xue, Q., Wang, Y.Q., Tsay, D., He, B., Jablons, D.M., and Rosen, S.D. (2010). Sulf-2, a heparan sulfate endosulfatase, promotes human lung carcinogenesis. Oncogene 29, 635–646.

Li, J.-P., and Kusche-Gullberg, M. (2016). Heparan Sulfate: Biosynthesis, Structure, and Function. Int. Rev. Cell Mol. Biol. 325, 215–273.

McAtee, C.O., Barycki, J.J., and Simpson, M.A. (2014). Emerging roles for hyaluronidase in cancer metastasis and therapy. Adv. Cancer Res. 123, 1–34.

Mead, T.J., McCulloch, D.R., Ho, J.C., Du, Y., Adams, S.M., Birk, D.E., and Apte, S.S. (2018). The metalloproteinase-proteoglycans ADAMTS7 and ADAMTS12 provide an innate, tendon-specific protective mechanism against heterotopic ossification. JCI Insight 3.

Morimoto-Tomita, M., Uchimura, K., Werb, Z., Hemmerich, S., and Rosen, S.D. (2002). Cloning and characterization of two extracellular heparin-degrading endosulfatases in mice and humans. J Biol Chem 277, 49175–49185.

Noborn, F., Gomez Toledo, A., Sihlbom, C., Lengqvist, J., Fries, E., Kjellen, L., Nilsson, J., and Larson, G. (2015). Identification of chondroitin sulfate linkage region glycopeptides reveals prohormones as a novel class of proteoglycans. Mol Cell Proteomics 14, 41–49.

Pasquato, A., Dettin, M., Basak, A., Gambaretto, R., Tonin, L., Seidah, N.G., and Di Bello, C. (2007). Heparin enhances the furin cleavage of HIV-1 gp160 peptides. FEBS Lett 581, 5807–5813.

Pempe, E.H., Burch, T.C., Law, C.J., and Liu, J. (2012). Substrate specificity of 6-O-endosulfatase (Sulf-2) and its implications in synthesizing anticoagulant heparan sulfate. Glycobiology 22, 1353–1362.

Pérard, J., Nader, S., Levert, M., Arnaud, L., Carpentier, P., Siebert, C., Blanquet, F., Cavazza, C., Renesto, P., Schneider, D., et al. (2018). Structural and functional studies of the metalloregulator Fur identify a promoter-binding mechanism and its role in Francisella tularensis virulence. Commun. Biol. 1, 93.

Peterson, S.M., Iskenderian, A., Cook, L., Romashko, A., Tobin, K., Jones, M., Norton, A., Gomez-Yafal, A., Heartlein, M.W., Concino, M.F., et al. (2010). Human Sulfatase 2 inhibits in vivo tumor growth of MDA-MB-231 human breast cancer xenografts. BMC Cancer 10, 427.

Petoukhov, M.V., Franke, D., Shkumatov, A.V., Tria, G., Kikhney, A.G., Gajda, M., Gorba, C., Mertens, H.D.T., Konarev, P.V., and Svergun, D.I. (2012). New developments in the ATSAS program package for small-angle scattering data analysis. J. Appl. Crystallogr. 45, 342–350.

Rosen, S.D., and Lemjabbar-Alaoui, H. (2010). Sulf-2: an extracellular modulator of cell signaling and a cancer target candidate. Expert Opin Ther Targets 14, 935–949.

Sarrazin, S., Lamanna, W.C., and Esko, J.D. (2011). Heparan sulfate proteoglycans. Cold Spring Harb Perspect Biol 3.

Seffouh, A., Milz, F., Przybylski, C., Laguri, C., Oosterhof, A., Bourcier, S., Sadir, R., Dutkowski, E., Daniel, R., van Kuppevelt, T.H., et al. (2013). HSulf sulfatases catalyze processive and oriented 6-O-desulfation of heparan sulfate that differentially regulates fibroblast growth factor activity. Faseb J 27, 2431–2439.

Seffouh, A., El Masri, R., Makshakova, O., Gout, E., Hassoun, Z.E.O., Andrieu, J.-P., Lortat-Jacob, H., and Vivès, R.R. (2019a). Expression and purification of recombinant extracellular sulfatase HSulf-2 allows deciphering of enzyme sub-domain coordinated role for the binding and 6-O-desulfation of heparan sulfate. Cell. Mol. Life Sci. CMLS 76, 1807–1819.

Seffouh, I., Przybylski, C., Seffouh, A., El Masri, R., Vivès, R.R., Gonnet, F., and Daniel, R. (2019b). Mass spectrometry analysis of the human endosulfatase Hsulf-2. Biochem. Biophys. Rep. 18, 100617.

Svergun, D.I. (1992). Determination of the regularization parameter in indirect-transform methods using perceptual criteria. J. Appl. Crystallogr. 25, 495–503.

Tang, R., and Rosen, S.D. (2009). Functional consequences of the subdomain organization of the sulfs. J Biol Chem 284, 21505–21514.

Tian, C., Öhlund, D., Rickelt, S., Lidström, T., Huang, Y., Hao, L., Zhao, R.T., Franklin, O., Bhatia, S.N., Tuveson, D.A., et al. (2020). Cancer Cell-Derived Matrisome Proteins Promote Metastasis in Pancreatic Ductal Adenocarcinoma. Cancer Res. 80, 1461–1474.

Uchimura, K., Morimoto-Tomita, M., Bistrup, A., Li, J., Lyon, M., Gallagher, J., Werb, Z., and Rosen, S.D. (2006). HSulf-2, an extracellular endoglucosamine-6-sulfatase, selectively mobilizes heparin-bound growth factors and chemokines: effects on VEGF, FGF-1, and SDF-1. BMC Biochem 7, 2.

Vives, R.R., Seffouh, A., and Lortat-Jacob, H. (2014). Post-Synthetic Regulation of HS Structure: The Yin and Yang of the Sulfs in Cancer. Front Oncol 3, 331.

Walhorn, V., Möller, A.-K., Bartz, C., Dierks, T., and Anselmetti, D. (2018). Exploring the Sulfatase 1 Catch Bond Free Energy Landscape using Jarzynski’s Equality. Sci. Rep. 8, 16849.

Yang, J.D., Sun, Z., Hu, C., Lai, J., Dove, R., Nakamura, I., Lee, J.S., Thorgeirsson, S.S., Kang, K.J., Chu, I.S., et al. (2011). Sulfatase 1 and sulfatase 2 in hepatocellular carcinoma: associated signaling pathways, tumor phenotypes, and survival. Genes. Chromosomes Cancer 50, 122–135.

Zhu, C., He, L., Zhou, X., Nie, X., and Gu, Y. (2016). Sulfatase 2 promotes breast cancer progression through regulating some tumor-related factors. Oncol. Rep. 35, 1318–1328.

